# Comparative analyses of saprotrophy in *Salisapilia sapeloensis* and diverse plant pathogenic oomycetes reveal lifestyle-specific gene expression

**DOI:** 10.1101/656496

**Authors:** Sophie de Vries, Jan de Vries, John M. Archibald, Claudio H. Slamovits

## Abstract

Oomycetes include many well-studied, devastating plant pathogens. Across oomycete diversity, plant-infecting lineages are interspersed by non-pathogenic ones. Unfortunately, our understanding of the evolution of lifestyle switches is hampered by a scarcity of data on the molecular biology of saprotrophic oomycetes, ecologically important primary colonizers of dead tissue that can serve as informative reference points for understanding the evolution of pathogens. Here, we established *Salisapilia sapeloensis* growing on axenic litter as a tractable system for the study of saprotrophic oomycetes. We generated multiple transcriptomes from *S. sapeloensis* and compared them to (a) 22 oomycete genomes and (b) the transcriptomes of eight pathogenic oomycetes grown under 13 conditions (three pathogenic lifestyles, six hosts/substrates, and four tissues). From these analyses we obtained a global perspective on the gene expression signatures of oomycete lifestyles. Our data reveal that oomycete saprotrophs and pathogens use generally similar molecular mechanisms for colonization, but exhibit distinct expression patterns. We identify *S. sapeloensis*’ specific array and expression of carbohydrate-active enzymes and regulatory differences in pathogenicity-associated factors, including the virulence factor EpiC2B. Further, *S. sapeloensis* was found to express only a small repertoire of effector genes. In conclusion, our analyses reveal lifestyle-specific gene regulatory signatures and suggest that, in addition to variation in gene content, shifts in gene regulatory networks might underpin the evolution of oomycete lifestyles.

## Introduction

Oomycetes encompass a great diversity of ecologically and economically relevant pathogens of plants [1,2]. These include, for example, the (in)famous *Phytophthora infestans*, the cause of the Irish potato famine [2]. In addition to the plant pathogenic genera other lineages are pathogens of animals [3,4]. Interspersed amongst the diversity of pathogenic oomycete lineages are (apparently) non-pathogenic lineages with saprotrophic lifestyles [5,6,7] (for an overview see Figure 1a). How saprotrophy arises in oomycetes is, however, not yet clear and depends strongly on the character of the last common ancestor of oomycetes. Two hypotheses have been considered: i) the last common ancestor was a pathogen and the saprotrophic lineages have arisen several times independently or ii) saprotrophy is the ancestral state in oomycetes [8].

**Figure 1.**
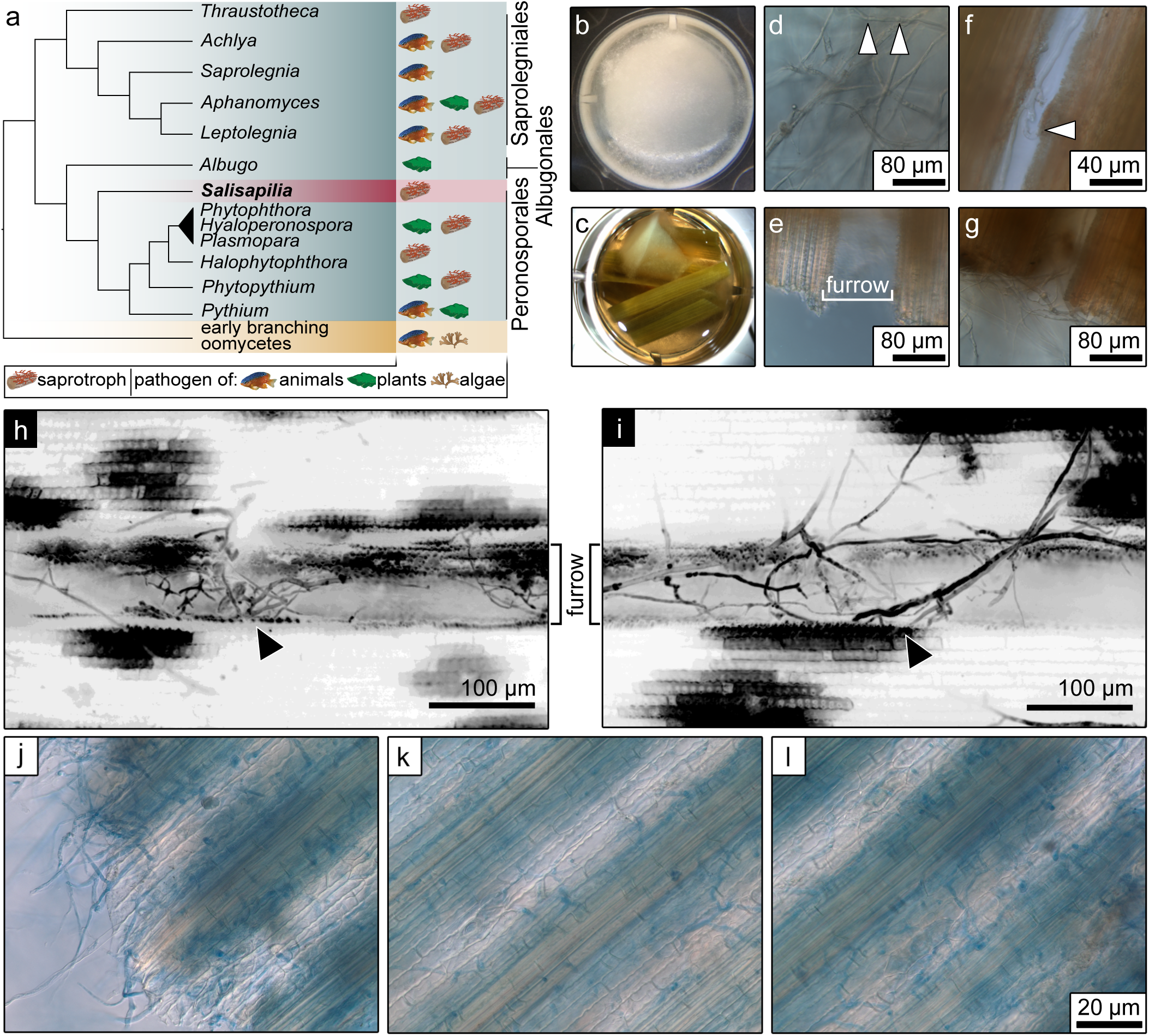
*Salisapilia sapeloensis*’ colonization of marsh grass litter. (a) Oomycetes phylogeny. The lifestyles that occur in a specific lineage are indicated on the right of each lineages. Lifestyles include: plant (leaf icon) and animal (fish icon) pathogens and saprotrophs (log icon) its occurrences are according to Diéguez-Uribeondo et al. [5], Beakes et al. [6], and Marano et al. [7]. The cladogram is based on published phylogenetic studies [6,24,110]. We note that based on more recent phylogenetic analyses the Salisapiliaceae, to which the genus *Salisapilia* belongs, could also be sister to the Peronosporaceae, which includes the polyphyletic genus *Phytophthora* [111], and would hence be embedded between *Phytophthora* and *Pythium*. (b,c) An established, axenic system for cultivating *S. sapeloensis*; closeups of *S. sapeloensis* growing in wells with liquid CMM (b) and MGL (c). (d-g) Micrographs of *S. sapeloensis*. (d) *S. sapeloensis* growing in water around the MGL. Septae are highlighted by arrow heads. (f) *S. sapeloensis* growing in the litter’s furrows and associated to leaf tissue (arrowhead). (e) Comparison of uninoculated and (g) inoculated MGL. (h,i) Calcofluor-white-stained (Fluorescent Brightener 28) *S. sapeloensis* growing into the furrows (labeled) and associating with leaf tissue (arrowheads; z-stacks reconstructed and projected from confocal micrographs; whole mount staining). It should be noted that calcofluor white stains both the cell walls of plants and oomycetes; more whole mount confocal micrographs can be found in Figure S1. (j-l) Micrographs of trypan blue stained leaf litter inoculated with *S. sapeloensis*.

Saprotrophic oomycetes are thought to play important ecological roles as colonizers and decomposers of organic matter, and by making organic debris more accessible to detritivores [7]. Furthermore, hybridization between saprotrophic and pathogenic *Phytophthora* spp. has been hypothesized to drive distant host jumps of this pathogenic lineage [7]. Additionally, the lifestyle of a microbe (pathogenic vs. non-pathogenic) depends on its environment, and potentially non-pathogenic organisms can be opportunistic pathogen under some environmental conditions [9,10,11,12]. Hence, any given saprotrophic oomycete lineage might in fact be a latent pathogen; we cannot exclude the possibility that they are able to colonize and exploit living plants. Despite the ecological functions of oomycete saprotrophs and their possible contributions to pathogenicity, how saprotrophs colonize their substrates and if and how they differ from pathogens remains little understood. Few genomic studies have begun to address this question.

Genomic data from *Thraustotheca clavata*, a saprotroph from the Saprolegniales, provided the first insights into the molecular biology of a saprotrophic oomycete [13]. The *in silico* secretome of *T. clavata* is similar in its predicted functions to the secretomes of oomycete pathogens. This is in agreement with genomic analyses of fungi showing that saprotrophs utilize a similar molecular toolkit as fungal pathogens [14,15]. This molecular toolkit includes, among other things, carbohydrate-active enzymes (CAZymes) for substrate degradation and cell wall remodelling, as well as other degradation-relevant enzymes such as peptidases and lipases, and sterol-binding elicitins [13,14,15]. Differences between pathogens and saprotrophs are, for example, found in the size of gene families, such as the cutinase family, which is required for successful pathogen infection but much smaller in numbers in the genomes of fungal saprotrophs [15]. The CAZyme content of fungi also seems to be shaped by lifestyles, hosts and substrates [16]. In a study of 156 fungal species, degradation profiles were found to be strongly substrate-dependent, although on similar substrates the ecological role (pathogen or saprotroph) was relevant [17].

One of the principal ideas in studying the genomics of filamentous microbes is that the history of the changes in their lifestyles can be inferred from their genomes [18,19]. Indeed, expansions or reductions of specific gene families can speak to the relevance of such genes for a certain lifestyle. Yet, gene presence / absence data does not provide insight into the *usage* of these genes by an organism with a given lifestyle. Moreover, as noted above, it cannot be ruled out that some saprotrophic oomycetes are latent pathogens—conclusions based on genomic data alone should be taken with caution. It is here that large-scale transcriptomic and proteomic analyses have the potential to provide deeper insight. Indeed, analysis of gene expression patterns is a powerful discovery tool for gaining global insight into candidate genes involved in distinct lifestyles, e.g., with virulence-associated and validated candidates being up-regulated during infection [20,21,22,23]. Such data underpin the notion that plant pathogenicity is at least partly regulated on the level of gene expression. Assuming that saprotrophic interactions are also at least partially regulated on the transcript level, we can study their molecular biology and differences relative to pathogens via gene expression analyses during a saprotrophic interaction.

Here we established the first axenic system for the study of saprotrophy in oomycetes. This system entailed the oomycete *Salisapilia sapeloensis* grown on sterilized marsh grass litter. Using this system, we applied a comparative transcriptomic approach to investigate molecular differences (a) within one saprotrophic interaction across three different conditions and (b) between the saprotrophic and several pathogenic interactions. Our data show *S. sapeloensis* in a successful saprotrophic interaction in which it actively degrades litter. These data are corroborated by the fact that CAZymes were found to be highly responsive in litter-associated and litter-colonizing mycelium. Comparing *S. sapeloensis* to plant pathogenic oomycetes, we identified transcriptomic signatures of saprotrophy and pathogencity. Our data point to differences in regulatory networks that may shape the evolution of oomycete lifestyles.

## Results and discussion

### *Salisapilia sapeloensis:* tractable saprotrophy under axenic lab conditions

Oomycetes have evolved both pathogenic as well as saprotrophic lifestyles. While many pathogenic oomycetes are being investigated in depth, little is known about the molecular basis of saprotrophic oomycetes. Comparative functional genomics of plant-microbe interactions in the context of different lifestyles can yield vital clues about what distinguishes the lifestyles of pathogens from those of non-pathogenic ones. To enable broad comparative functional genomic investigations of oomycete biology, we are studying *Salisapilia sapeloensis* for three main reasons: i) it is a peronosporalean oomycete, ii) it has an interesting phylogenetic position basal to many plant pathogenic oomycetes [24] (see also Figure 1a) and iii) its natural substrate is known and easily accessible.

*S. sapeloensis* was originally isolated from the litter of the marsh grass *Spartinia alterniflora* [24]. To study *S. sapeloensis* in the lab, we created axenic marsh grass litter (MGL). We then inoculated corn meal medium (CMM, control) as well as MGL submersed in salt water with mycelium of *S. sapeloensis* (Figure 1b,c).

The mycelium of *S. sapeloensis* grew well in both medium and the salt water next to the litter (Figure 1b-d). To study if and how *S. sapeloensis* associated with the litter, we investigated the inoculated MGL microscopically. *S. sapeloensis* was frequently observed around the edges of axenic MGL (Figure 1e,g) and grew into its furrows (Figure 1f,h,i, Figure S1). The mycelium associated closely with leaf tissue in the furrows (Figure 1f) and was also observed growing inside the tissue (Figure 1j,k,l). Additionally, we noted that the water surrounding the inoculated tissue turned yellow in color (Figure 2a)—likely a result of litter degradation, further supporting the notion that *S. sapeloensis* utilizes axenic MGL as a substrate. Overall, *S. sapeloensis* thrived on the litter. Within seven days it produced enough biomass for our downstream analyses. Hence, *S. sapeloensis* is capable of colonizing and surviving on MGL, as the MGL contained no additional nutrients but those encased in the tissue of the marsh grass leaves.

**Figure 2.**
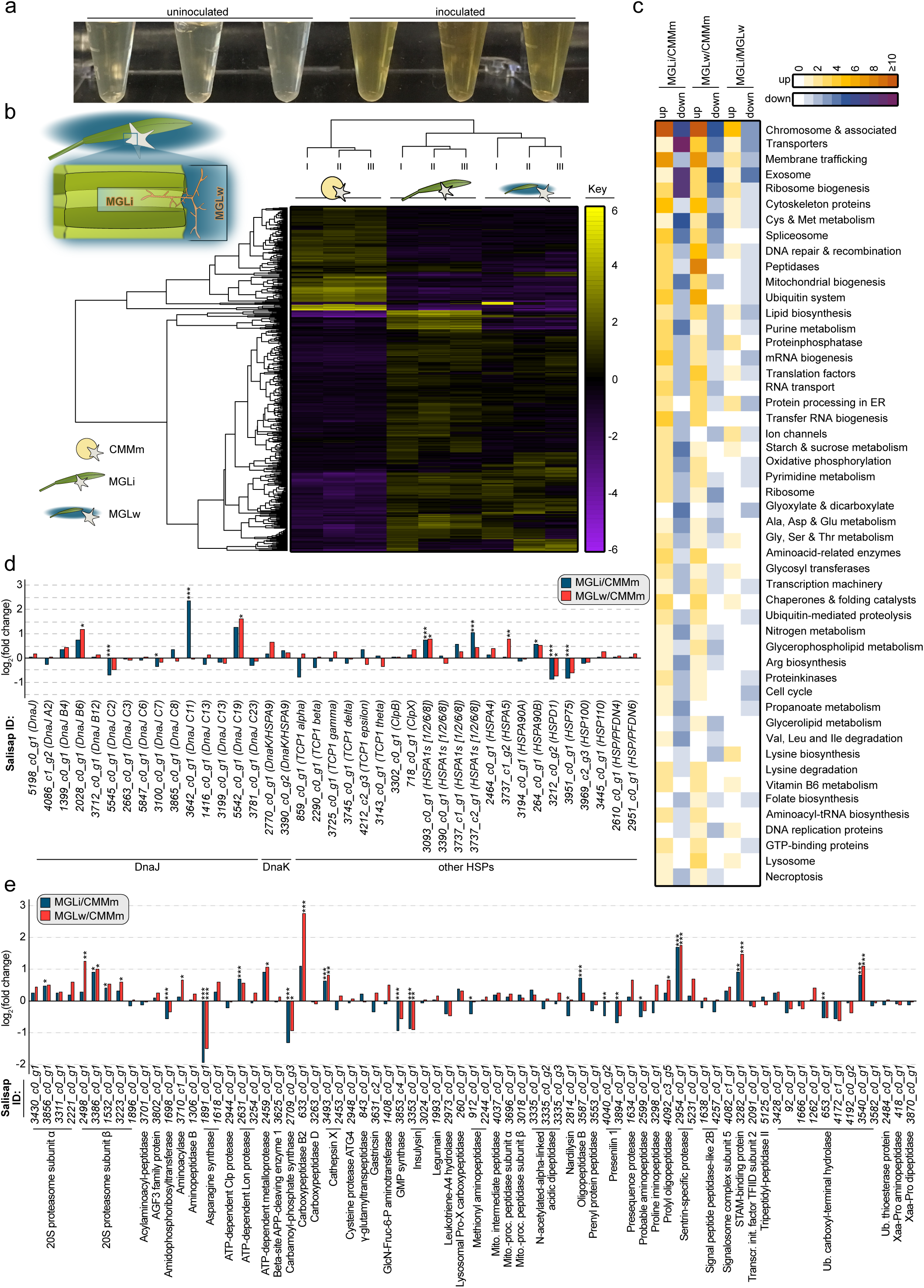
Transcriptomic profiling of the axenic saprotrophic system *Salisapilia sapeloensis* during colonization of litter. (a) Visual differences in the water next to uninoculated litter and litter inoculated with *S. sapeloensis*. (b) Left: Sampling strategy for MGLi and MGLw; right: hierarchically clustered expression values (log_2_-transformed; median-centered) for CMMm, MGLw and MGLi. Higher expression than the median-centred expression is indicated in yellow, lower expression is indicated in purple. I, II and III indicates the biological replicates for each condition. (c) The 50 most responsive KEGG pathways/BRITE hierachies, indicating the overall number of KEGG orthologs in a given KEGG pathway/BRITE hierachy with an induced (white to orange) or reduced (white to dark purple) expression. (d) log_2_(FC) of the genes corresponding to KEGG orthologs in the umbrella category ‘Chaperone and folding catalysts’. Note that one KEGG ortholog can include more than one gene. Only genes with a TPM_TMM-normalized_≥1 in at least one of the three conditions are shown. The y-axis indicates the log_2_(FC) when MGLi (blue) or MGLw (red) is compared to CMMm. The gene IDs are given below the bar graph for each gene and in brackets the corresponding KEGG ortholog is indicated. Overall functional categories are given below the gene IDs. Significant differences between MGLi or MGLw vs. CMMm are indicated by * p-value ≤ 0.05, ** p-value ≤ 0.01 and *** p-value ≤ 0.001. (e) log_2_(FC) of the genes corresponding to KEGG orthologs in the umbrella category ‘Peptidases’. The figure shows only genes with a TPM_TMM-normalized_ ≥ 1 in at least one treatment. The comparison MGLi vs. CMMm is shown in blue and MGLw vs. CMMm is shown in red. The gene IDs are indicated below the bars. Below the gene IDs we noted the corresponding KEGG orthologs. Significant differences are given as * FDR ≤ 0.05, ** FDR ≤ 0.01 and *** FDR ≤ 0.001.

### Molecular signatures of litter colonization

To more fully elucidate how *S. sapeloensis* colonizes and lives off litter, we isolated RNA in biological triplicates and performed RNAseq on cultures growing under three conditions: *S. sapeloensis* grown in regular CMM medium (henceforth called CMMm), in water next to MGL (MGLw) and on MGL (MGLi; Figure S2). *De novo* assembly resulted in 7777 unique genes (Dataset S1a). Of those, 4592 were oomycete-affiliated and 2656 were orphans (Dataset S1b,c); the remaining 529 genes had a taxonomic affiliation to other organisms and have hence been omitted from downstream analyses. Unless otherwise mentioned, downstream analyses focused on the strict oomycete-affiliated dataset. Overall, 1628 oomycete-affiliated genes were differentially expressed in at least one comparison (MGLi vs. CMMm, MGLi vs. MGLw and MGLw vs. CMMm; Benjamini-Hochberg adjusted *P* [FDR]≤0.05; Dataset S1d). Of those 1628 genes with differential expression, 90% (1472 genes) are found in the pairwise comparison of MGLi vs. CMMm. In agreement with different sets of biochemical cues delivered by the two nutritional resources in our growth conditions, the expression profiles of MGLi were more similar to MGLw than to CMMm (Figure 2b). Nonetheless, 145 oomycete-affiliated genes showed differential expression patterns (FDR≤0.05; Dataset S1d) between MGLi and MGLw. These are, therefore, candidate genes specifically associated with the colonization and on-site degradation of the litter.

Next, we determined which pathways the saprotroph requires for litter colonization. To do so, we classified the assembled genes using the KEGG pathways as well as BRITE functional hierarchies (Dataset S2a); we then determined which umbrella categories in that classification were most responsive in *S. sapeloensis* (Figure 2c). Here we define ‘responsiveness’ as the number of KEGG orthologs with a cumulative 2-fold change (log_2_ (fold change [FC]) ≥1 or ≤−1) in a given umbrella category in pairwise comparisons of MGLi, MGLw and CMMm.

Among the 50 most responsive KEGG pathways/BRITE hierarchy terms, three showed exclusively an induction (log_2_ FC ≥1) upon exposure to MGL. These are ‘Chaperone and folding catalysts’, ‘Vitamin B6 metabolism’, and ‘Lysosome’ (Figure 2c). ‘Chaperone and folding catalysts’ suggests an association with stress: Of the 48 KEGG orthologs present in the ‘Chaperone and folding catalysts’ pathway, 34 belonged to the heat shock protein family: 12 were annotated as DnaJ KEGG orthologs, one was annotated as a DnaK KEGG ortholog and 19 were annotated as other heat-shock protein (HSP) KEGG orthologs. This corresponds to 15 DnaJ-encoding, two DnaK (HSPA09)-encoding and 26 HSP-encoding genes (Dataset S2b). All except four HSP-encoding genes had a TPM_TMM-normalized_ ≥1, i.e., 39 genes were sufficiently expressed. In this dataset seven of the 39 genes are significantly higher expressed in litter-associated treatments (i.e., MGLi or MGLw vs. CMMm; Figure 2d). In a study by Benz et al. [25], transcripts for stress-associated proteins—such as DnaK—showed an up-regulation in the saprotrophic fungus *Neurospora crassa* when exposed to different plant cell wall-associated carbon sources. Hence, our data suggest that *S. sapeloensis* experiences stress when colonizing and degrading the plant tissue as well. It is conceivable that toxic degradants and stored plant compounds trigger such stress in the saprotrophic oomycete. Alternatively, this observation could be the result of secretion stress, which has also been observed in saprotrophic fungi [26]. This hypothesis is supported by the responsiveness of other KEGG pathways/BRITE hierarchy terms, such as ‘Protein processing in the ER’ or ‘Membrane trafficking’ (Figure 2c).

Vitamin B6 is a precursor of pyridoxal phosphate, a co-factor for protein synthesis [27]. Hence, the elevation of the KEGG pathway ‘Vitamin B6 metabolism’ in *S. sapeloensis* exposed to MGL might be related to protein degradation and protein biosynthesis. This is in agreement with the presence of several amino acid synthesis-related KEGG pathways/BRITE hierarchy terms among the 50 most responsive categories (Figure 2c). An elevation of the KEGG-pathway “Lysosome” further indicates an increase in degradation of biological material, in agreement with a saprotrophic lifestyle. Given that stress-associated genes also are up-regulated during saprotrophy in fungi [25], the increase of the BRITE term ‘Chaperone and folding catalysts’, and the KEGG-pathways ‘Vitamin B6 metabolism’ and ‘Lysosome’ in MGLi or MGLw vs. CMMm indicates a general change in functional degradation in *S. sapeloensis*. We therefore investigated the direction of responsiveness in KEGG-pathways associated with degradation in more detail.

The MGLw and MGLi samples contained no nutritional carbon source other than the grass litter. In this situation, *S. sapeloensis* thus had to live off the litter. Our macro- and microscopic investigations (Figure 1b-l, 2a) suggest that *S. sapeloensis* is capable of doing so. This observation is now underpinned by the responsiveness of the KEGG pathways ‘Lysosome’ and ‘Starch and sucrose metabolism’, and the BRITE terms ‘Transporters’, ‘Membrane trafficking’, and ‘Peptidases’ (Figure 2c). Both ‘Transporters’ and ‘Membrane trafficking’ are among the three most responsive KEGG pathways/BRITE hierarchy terms. Both have KEGG orthologs induced and reduced in transcript abundance in the pairwise comparisons. Their responsiveness hence speaks not just to a mere induction but to specific differential up- and down-regulation as well. ‘Membrane trafficking’ contains 164 KEGG orthologs of which 14 responded to the growth condition: 10 KEGG orthologs, corresponding to 10 genes, were increased in MGLi and MGLw vs. CMMm (log_2_(FC) ≥1, Dataset S2), while four KEGG orthologs, corresponding to four genes, were reduced in MGLi and MGLw vs. CMMm (log_2_(FC) ≤−1, Dataset S2). Therefore, we observed an overall tendency for an induction of ‘Membrane trafficking’. Indeed, in fungi the responsiveness of the secretory pathway components is correlated with the degradation of plant material [25,28]. Such connections are also likely to exist in oomycetes. Moreover, both visual clues (Figure 2a) and the responsiveness of degradation-associated KEGG pathways/BRITE hierarchy terms provide evidence of *S. sapeloensis* actively degrading the litter substrate.

We next focused on the degradation-associated responsive categories, i.e., ‘Peptidase’ and ‘starch and sucrose metabolism’. The BRITE term ‘Peptidase’ included 77 KEGG orthologs, 73 of which had an average TPM_TMM-normalized_ ≥1 (Figure 2e). ‘Peptidase’ is induced in MGLw vs. CMMm, and more KEGG orthologs have an induced transcript abundance than a reduced transcript abundance when MGLi is compared to CMMm (Figure 2c). 27 peptidase-encoding genes were differentially expressed in one of the three comparisons (FRD≤0.05). Of those 27 peptidase-encoding genes, 56% (i.e., 15 genes) are up-regulated in either MGLi or MGLw vs. CMMm (Figure 2e). In agreement with the KEGG ortholog analyses, the majority of the up-regulated peptidase-encoding genes is further attributable to the comparison of MGLw against CMMm. This suggests that mycelium growing next to the litter majorly contributes to the degradation of proteins in *S. sapeloensis*.

The alterations in gene expression in the KEGG pathway ‘starch and sucrose metabolism’ suggest a remodelling of carbon-based degradation upon the change of substrate. Such remodelling is expected, because CMMm, MGLi and MGLw represent different carbon sources, with CMMm, for example, being enriched in starch compared to MGL or the salt water. Likewise, MGL will have long carbon-chains, such as cellulose, which might reach the salt water in an already pre-processed condition. The molecular differences observed when we dissect the components of this pathway align with the differences in carbon sources: For example, the KEGG ortholog endo-1,4-β-D-glucanase (EC 3.2.1.4), composed of 10 endo-1,4-β-D-glucanase-encoding genes, is overall induced (log_2_ [FC] >1, Dataset S2) in MGLi vs. MGLw and CMMm. Three of the 10 endo-1,4-β-D-glucanase-encoding genes contribute to both the induction in MGLi vs. MGLw and MGLi vs. CMMm (*Salisap3873_c0_g6, Salisap3873_c0_g7* and *Salisap3873_c0_g8*; Dataset S1d). Endo-1,4-β-D-glucanase is involved in the degradation of glucans found in leaves [29], highlighting the ability of *S. sapeloensis* to degrade leaf litter. This is further illustrated by the extent to which the three endo-1,4-β-D-glucanase-encoding genes that caused the overall induction were up-regulated: All three genes were on average 87-fold up-regulated (FDR≤0.05) in MGLi vs. CMMm and 49-fold up-regulated (FDR≤0.05) in MGLi vs. MGLw. One of the three genes (*Salisap3873_c0_g8*) even increased by 248-fold in MGLi vs. CMMm (FDR = 5.8*10^−46^) and 109-fold in MGLi vs. MGLw (FDR = 5.3*10^−33^).

In contrast to the overall induction of endo-1,4-β-D-glucanase, the KEGG ortholog phosphoglucomutase (represented by one gene in the database, *Salisap4157_c0_g2*) has a reduced transcript abundance in MGLi vs CMMm (log_2_(FC)<-1; Dataset S2a). This is substantiated by the differential gene expression analysis, showing that the single phosphoglucomutase-encoding gene is significantly down-regulated by 2-fold (FDR = 5.7*10^−10^) in MGLi vs. CMMm. Phosphoglucomutase is associated with starch catabolism [30,31], illustrating the reduced availability of starch in litter compared to corn meal-based medium. Taken together, the visual and molecular data show that *S. sapeloensis* acts as a saprotroph in our axenic set-up—making it an apt system for studying oomycete saprotrophy *in vitro*.

Notably, all four degradation-associated categories had KEGG orthologs specifically responsive (log2 FC>1/<-1) in MGLi vs MGLw (Figure 2c). This suggests that MGLw has a slightly different metabolism than MGLi. This makes sense given that the water next to the litter presumably contains pre-processed organic compounds. We therefore interpret this different regulation of biochemical and metabolic pathways as a division of labor, in which mycelium growing on litter (i.e. proximal) mainly degrades large carbon-components into smaller pieces that are further processed by mycelium grown in the surrounding water (i.e. distal). Such division of labor has been observed across different mycelial parts in *Sclerotinia sclerotiorum*, an ascomycete growing as a septate mycelium [32]. Peyraud and colleagues [32] hypothesized that cooperation of distinct mycelial parts in a multicellular mycelium may be of an advantage to the pathogen. Indeed, their study indicated that the division in metabolic tasks would improve plant colonization. Given that the mycelium of *S. sapeloensis* is also septate [24] (see also Figure 1d), such division of labor is likewise feasible. Similar to *S. sclerotiorum*, the two distinct parts of *S. sapeloensis* show significant differences with regard to the degradation of plant material. Hence, even though *S. sapeloensis* acts as a saprotroph in our system, it is capable of division of labor, which may give it an advantage in colonizing the dead plant tissue.

### *S. sapeloensis* expresses a distinct degradation enzyme profile

The transcriptomic ‘fingerprints’ of *S. sapeloensis* show that the saprotroph actively degrades MGL. Degradation of plant material is, however, not restricted to saprotrophy, but is also essential for plant pathogenicity [29]. To analyse if and how degradation processes differ between *S. sapeloensis* and oomycete pathogens, we first explored putative orthologous genes between *S. sapeloensis* and 16 peronosporalean plant pathogens and second analyzed those genes unique to *S. sapeloensis*. We identified on average 1376 genes in *S. sapeloensis* with no detectable ortholog in at least one pathogen; 705 of these genes (51.2%) are entirely unique to *S. sapeloensis* (Dataset S3). Among these unique genes, 191 (27% of 705) have significantly different expression (FDR≤0.05) in MGLi compared to CMMm (Dataset S3c). Similarly, 22% (153 genes) were differentially regulated in MGLw vs. CMMm (FDR≤0.05), but only 31 (4.3%) are differentially expressed in MGLi vs. MGLw. To identify the gross molecular functions of the differentially regulated, unique genes we performed a Gene Ontology (GO-)enrichment analysis for the categories MGLi vs CMMm and MGLw vs. CMMm. This analysis pointed to a specific enrichment in hydrolase activity (Figure 3a), and implies that *S. sapeloensis* has a novel profile of degradation enzyme compared to the oomycete plant pathogens.

**Figure 3.**
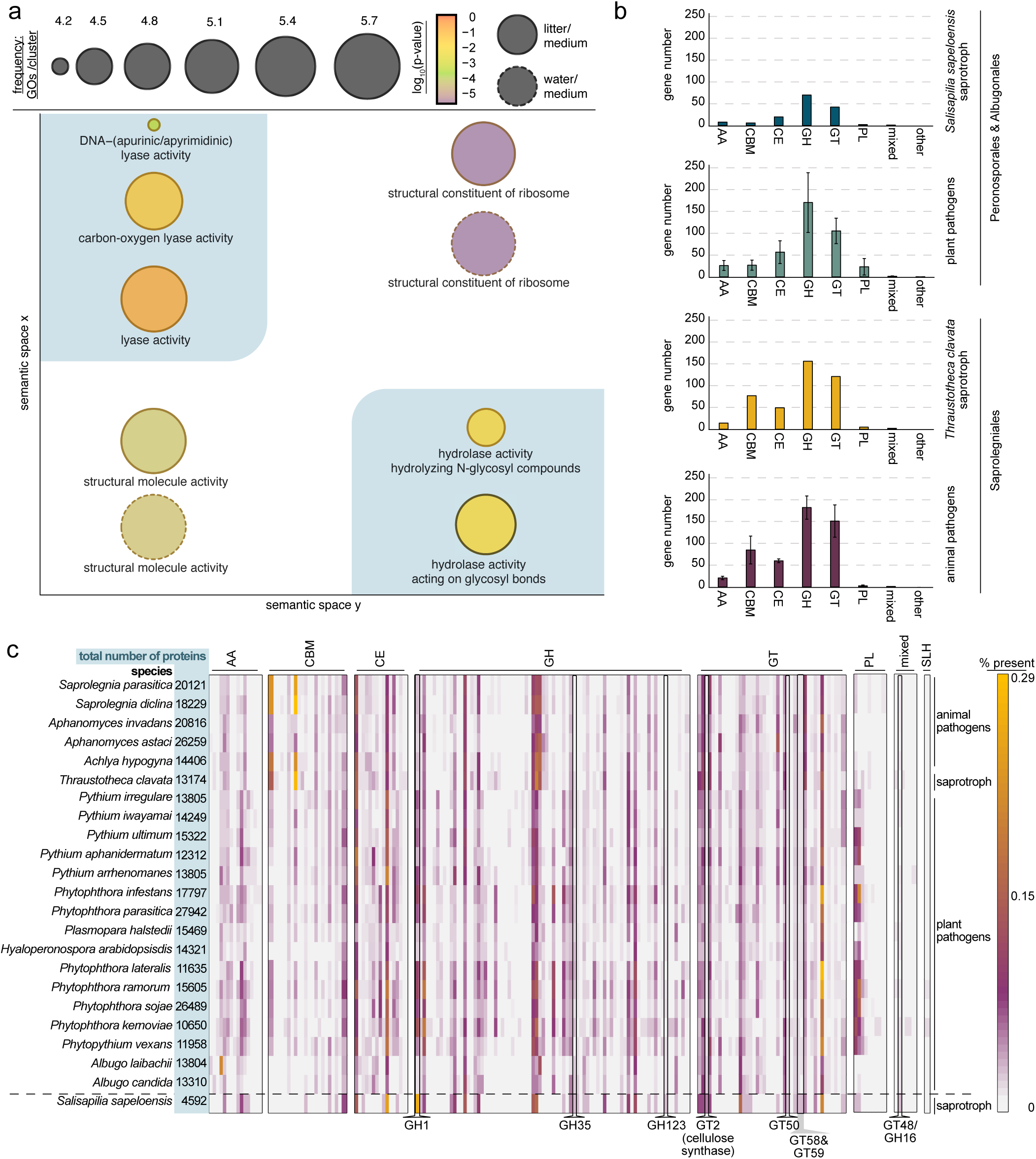
Comparative analyses of the CAZyme repertoire of *Salisapilia sapeloensis*. (a) GO-term enrichment of genes unique to *S. sapeloensis*, which are up-regulated in MGLi/MGLw vs. CMMm (GO-term frequency=circle size, log_2_(p-value) is indicated by the colors in the circle). The solid lines around the circles indicate an enrichment in MGLi/CMMm and the dotted lines around a circle indicate an enrichment in MGLw/CMMm. The blue background highlights lyase and hydrolase activity. (b) CAZyme distribution across oomycetes. (c) Heatmap showing the relative number of CAZymes in relation to the overall number of all analysed *in silico* translated proteins (%) in an oomycete species (total number of proteins is given to the right of each species name). The 23 oomycete species are noted on the left; their lifestyles are noted on the right. Every vertical line represents a CAZyme family. Highlighted are CAZyme-encoding gene families with an increased relative presence (i.e. log_2_(FC)≥1) in *S. sapeloensis*’ transcriptome vs. oomycete genomes. These are hence gene families that are expressed more than expected. The relative presence is indicated from white = 0% to yellow = 0.29%.

To further investigate the mechanisms underlying the degradation of litter and how they differ to those involved in the degradation of living plant material, we predicted the functional composition of the CAZyme repertoire of *S. sapeloensis*. *S. sapeloensis* expressed in total 149 genes predicted to encode CAZymes (134 with TPM_TMM-normalized≥_1 in at least one treatment; Dataset S4). Given that our dataset is a transcriptome, *S. sapeloensis* likely harbors more CAZyme-encoding genes in its genome than it expresses. However, the relative representation of CAZyme-encoding genes in the transcriptome of *S. sapeloensis* can give us some insights into their relevance for its saprotrophic lifestyle in our *in vitro* system. To get a measure on this relative representation (in absence of genomic data from *S. sapeloensis*), we analysed 22 genomes from diverse oomycetes to estimate the average amount of CAZyme-encoding genes.

We detected between 242 (*Albugo candida*, 1.8% of the protein-coding repertoire) and 731 (*Phytophthora parasitica*, 2.6% of the protein-coding repertoire) CAZyme-encoding genes in the oomycete genomes (Dataset S4a). On average, the oomycete genomes harboured 431±139 CAZyme-encoding genes. Assuming a similar amount for the saprotroph, *S. sapeloensis* expresses 38.2%±12.2% of the average CAZyme repertoire. The distribution of numbers of CAZyme-encoding genes detected in the *S. sapeloensis* transcriptome was similar to that encoded in genomes of plantpathogenic oomycetes, with the exception of polysaccharide lyases (PLs) (Figure 3b). This reduced representation of PLs in the saprotroph (relative to the plant pathogenic oomycetes) may be because of its substrate: Monocot cell walls have lower pectin content than their dicot counterparts [33]. The most abundant PLs in oomycetes (PL1 and PL3; Dataset S4a) are involved in pectin and pectate degradation [29] and most plant pathogens in our dataset infect dicots. *S. sapeloensis* in contrast colonizes a monocot, explaining the low number of PLs expressed. Therefore, *S. sapeloensis* expresses an overall representative number of CAZyme-encoding genes.

In contrast to the Peronosporales and Albugonales, all Saprolegniales (regardless of trophism) showed an increase in carbon binding module (CBM)-containing proteins (Figure 3b). Overall, the CAZyme repertoire of *S. sapeloensis* is more similar to that of the other plant pathogens than to the saprolegnian saprotroph. The representation of the six major CAZyme categories in a genome is, therefore, rather determined by phylogenetic position than lifestyle (Figure 1a). Nevertheless, our GO-term analyses showed that *S. sapeloensis*’ unique genes were associated with CAZyme-encoding genes. While, at this point, we cannot exclude that these gains are lineage-specific to *S. sapeloensis*, we observe that this picture mirrors what we know from fungi, where the lifestyle does shape the CAZyme repertoire [16]. We hence suspected that the differences in the CAZyme repertoire might be found in specific families, to which we next turned our attention.

### Pathogenicity-associated CAZyme families are not expressed during the saprotrophic interaction

In peronosporalean plant pathogens, CAZymes in general and glycosyl hydrolases (GHs) in particular, constitute a large fraction of the genes that were suggested to have been horizontally acquired from fungi, thus furthering peronosporalean plant pathogenicity [34]. *S. sapeloensis*, whose phylogenetic placement is basal to the peronosporalean plant pathogens (Figure 1a), does not express these particular GH families in any of the here tested conditions (Dataset 4), supporting their relevance for plant pathogenicity. This hints that *S. sapeloensis* does not express pathogenicity-associated CAZyme families during a saprotrophic interaction. To substantiate this finding, we next mined *S. sapeloensis*’ transcriptome for those CAZyme families that are induced during infection, using *Phytophthora infestans* as the reference point. Induced CAZyme families during infection of potato tubers by *P. infestans* are Carbohydrate esterase 8 (CE8), GH28, GH53, GH78, PL1, PL2 and PL3 [35]. Of these seven families only two (PL1 and PL3) were found to be expressed in *S. sapeloensis* (Dataset S4a). Hence, as we hypothesized based on the GO-term enrichment analyses (Figure 3a), the saprotroph seems to use different CAZyme families compared to plant pathogens.

To highlight CAZymes families with specific relevance for saprotrophy, we determined which genes were up-regulated during litter colonization. *S. sapeloensis* expressed mainly GHs, glycosyltransferases (GTs) and CEs (Figure 3b,c). In agreement with litter degradation, 19.0% of GH-encoding and 27.8% of CE-encoding genes showed a TPM_TMM-normalized_>100 in MGLi (Figure 4a, Dataset S4c). Overall, 37 CAZyme-encoding genes were up-regulated in litter-associated growth vs. growth on CMMm (FDR≤0.05). We analysed the relative expression of four of those genes (*Salisap4165_c1_g3* [*CE10*]*, Salisap3316_c0_g4* [*GH1*]*, Salisap3873_c0_g7* [*GH6*] and *Salisap1930_c0_g1* [*GT57*]) using qRT-PCR (Figure 4b). The up-regulation in MGLi vs CMMm in all four genes was confirmed for three of them (*Salisap4165_c1_g3* [*CE10*]*, Salisap3316_c0_g4* [*GH1*] and *Salisap3873_c0_g7* [*GH6*]). *Salisap1930_c0_g1 (GT57)* was the only gene up-regulated in MGLw vs. CMMm in the transcriptomic data; this was likewise confirmed. One of the most expressed genes in the entire CAZyme dataset was *Salisap3873_c0_g7*, one of the three *GH6* that we already noted in the KEGG analyses*. Salisap3873_c0_g7* was not only up-regulated in MGLi vs. CMMm (FDR = 4.8*10^−15^) but also MGLi vs. MGLw (FDR = 1*10^−5^; Figure 4a,b, Dataset S1d), further supporting its relevance in litter degradation. Across all CAZyme categories, 7.6% (10 of 131 genes with a calculatable log_2_[FC]) were specifically up-regulated in MGLi vs. MGLw; only 2.3% (3 of 131 genes) were down-regulated (Figure 4c). Together this amounts to 8.9% of the genes that are differentially regulated genes in MGLi vs. MGLw. These data underpin the importance of differential regulation of CAZyme-encoding genes for—at least our case of—oomycete saprotrophy. Among these litter-specific up-regulated genes in *S. sapeloensis* are two CEs, six GHs and all PLs. Taken together, these results suggest that PLs, GHs and CEs play important roles in litter degradation.

**Figure 4.**
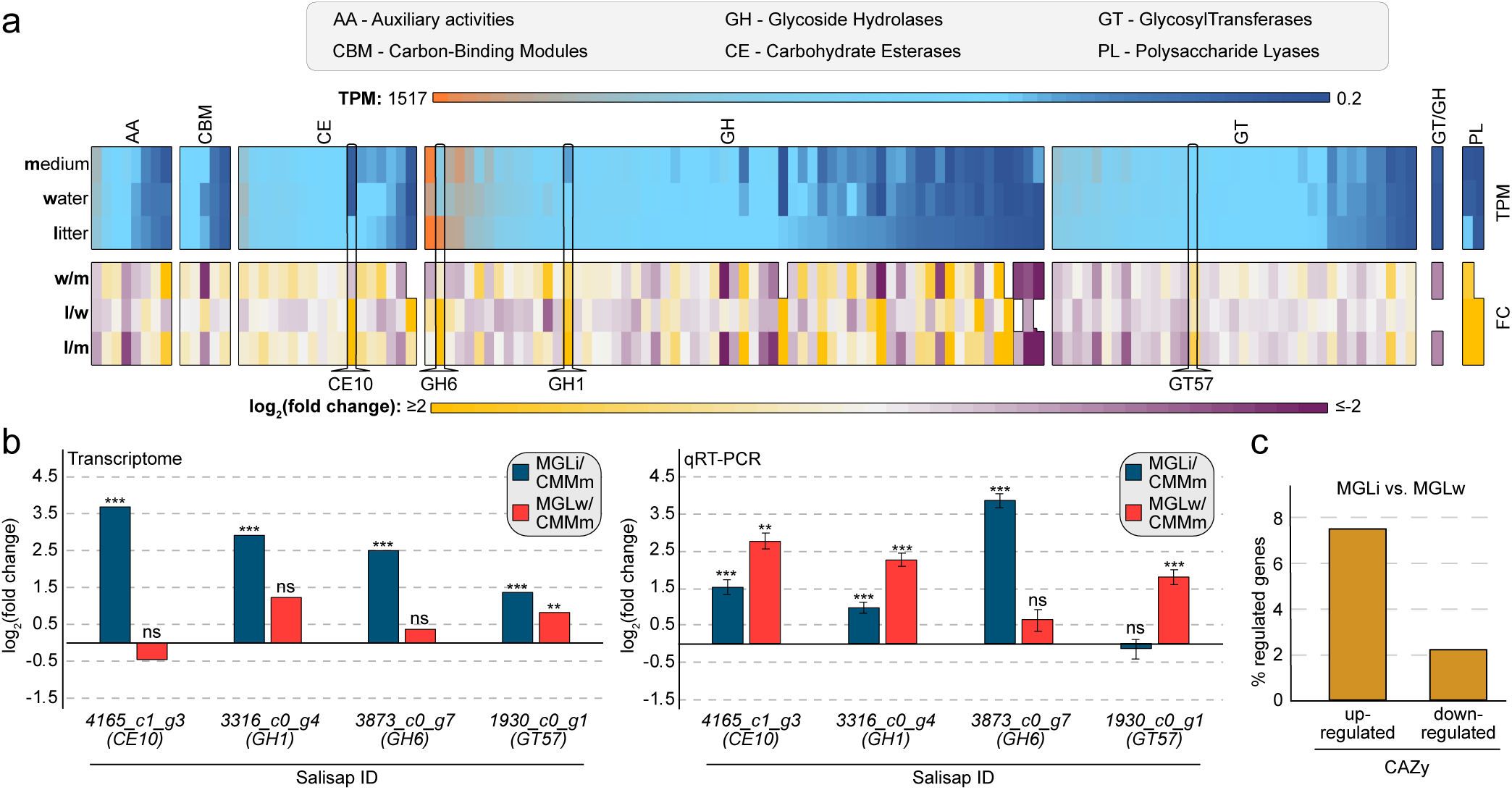
Expression of CAZyme-encoding genes in *Salisapilia sapeloensis*. (a) Expression of CAZyme-encoding genes in *S. sapeloensis*. Top: Heatmap of TPM_TMMnormalized_ values. The heatmap is sorted according to the expression in litter from highest (orange) to lowest (dark blue) in each CAZyme category. Medium indicates the expression in CMMm, water in MGLw and litter in MGLi. Each vertical line corresponds to one gene. Bottom: Heatmap of the log_2_(FC) in w/m (MGLw/CMMm), l/w (MGLi/MGLw) and l/m (MGLi/CMMm). The order of genes corresponds to the order of genes in the heatmap for TPM_TMMnormalized_ values on top. Four genes, which were further tested using qRT-PCR and mentioned in the main text, are highlighted. (b) Average relative expression (log_2_[FC]) of four genes up-regulated in MGLi and/or MGLw vs. CMMm in the transcriptome (left) and tested using qRT-PCR (right). The reference gene for the qRT-PCR was *SsH2A*. The error bars indicate the standard error of the mean (SEM). Gene IDs of the tested genes are given below the bar graphs. Significant differences are given above the bars as asterisk: * FDR/p-value ≤ 0.05, ** FDR/p-value ≤ 0.01, *** FDR/p-value ≤ 0.001. (c) Overall up- and down-regulation of CAZyme-encoding genes in litter colonization (MGLi vs. MGLw).

To find additional CAZyme families with a relevance in oomycete saprotrophy, we analyzed if and which CAZyme-encoding gene families have an enriched expression in *S. sapeloensis*. We reasoned that given the similar distribution pattern of CAZyme categories in the transcriptome and the oomycete genomes (Figure 3b), the relative amount of any CAZyme family specifically required for saprotrophy would be enhanced compared to the average relative representation in oomycete genomes. To that end we calculated the percentage of CAZyme-encoding genes in the transcriptome of *S. sapeloensis* and in the genomes of the other oomycetes (Figure 3c, Dataset S4b). We defined enrichment of a CAZyme-encoding family in the transcriptome as an at least two-fold increase in relative abundance of this family compared to its average abundance in the oomycete genomes (averaged over 22 genomes). Of the 191 CAZyme families detected, eight were more abundantly expressed than expected (Figure 3c). These eight families correspond to 14 GH-, five GT- and one mixed GH/GT-encoding gene (Figure 3c). Of these, seven were up-regulated in association with litter (MGLi and MGLw vs. CMMm; FDR ≤0.05; Dataset S1d). Six induced genes corresponded to the GH1 family (*Salisap3316_c0_g4*, *Salisap3316_c0_g3*, *Salisap3316_c0_g1*, *Salisap3477_c0_g1*, *Salisap3477_c0_g3* and *Salisap3920_c3_g1*) and one belonged to GT58 (*Salisap 2235_c0_g1*). Only one gene, a *GH1* (*Salisap3316_c0_g4*) is expressed specific to the colonization of litter, showing a 3.2-fold up-regulation (FDR=0.026; Dataset 1b). The GH1 family is generally involved in cellulose-degradation [29], highlighting the GH1 family, and especially *Salisap3316_c0_g4*, as a candidate CAZyme important for the saprotrophic lifestyle.

In summary, a large fraction of *S. sapeloensis*’ differentially regulated genes (i.e. FDR ≤0.05) in MGLi vs. MGLw encoded CAZymes. Additionally, CAZyme-encoding genes were highly responsive (strong changes in transcript levels and differential regulation) in comparisons between litter-associated growth and growth on CMM. This can be attributed to the changes in carbon sources that are degraded in MGLi, MGLw and CMMm. Likewise, CAZyme-encoding genes of fungi contribute a major fraction to the differential transcriptome after changes in carbon sources [25,36]. Additionally, our data highlights a distinct profile of CAZyme-encoding genes that are used during saprotrophy compared to plant pathogenicity. Taken together, this suggests that fungi and oomycetes co-opt their molecular biology in similar ways to similar lifestyles. Saprotrophy may hence have convergently evolved similar degradation mechanisms in these two groups of organisms.

### *Salisapilia sapeloensis*’ expression profile during litter decomposition is distinct from that of two plant pathogenic relatives during infection

We next sought to identify other hallmark genes of oomycetes associated with saprotrophy as opposed to plant pathogenicity. For this, we compared the global gene expression data from *S. sapeloensis* in MGLi vs. CMMm with published transcriptome-derived expression data from (i) *Phytophthora infestans* and (ii) *Pythium ultimum* infecting potato tubers vs. CMMm [23] (Figure 5). *S. sapeloensis* is ideal for this comparison because of its phylogenetic position basal to *Phytophthora* and *Pythium* [24] (Figure 1a). Moreover, the two pathogens have different modes of a pathogenic lifestyle: *P. infestans* is a hemibiotrophic pathogen. It requires a living host in its early infection stage (biotrophic phase) and later induces host cell death to use the degradants as a nutrient source (necrotrophic phase). In contrast, *P. ultimum* is a necrotrophic pathogen. It immediately kills its host after infection to utilize the degradants.

**Figure 5.**
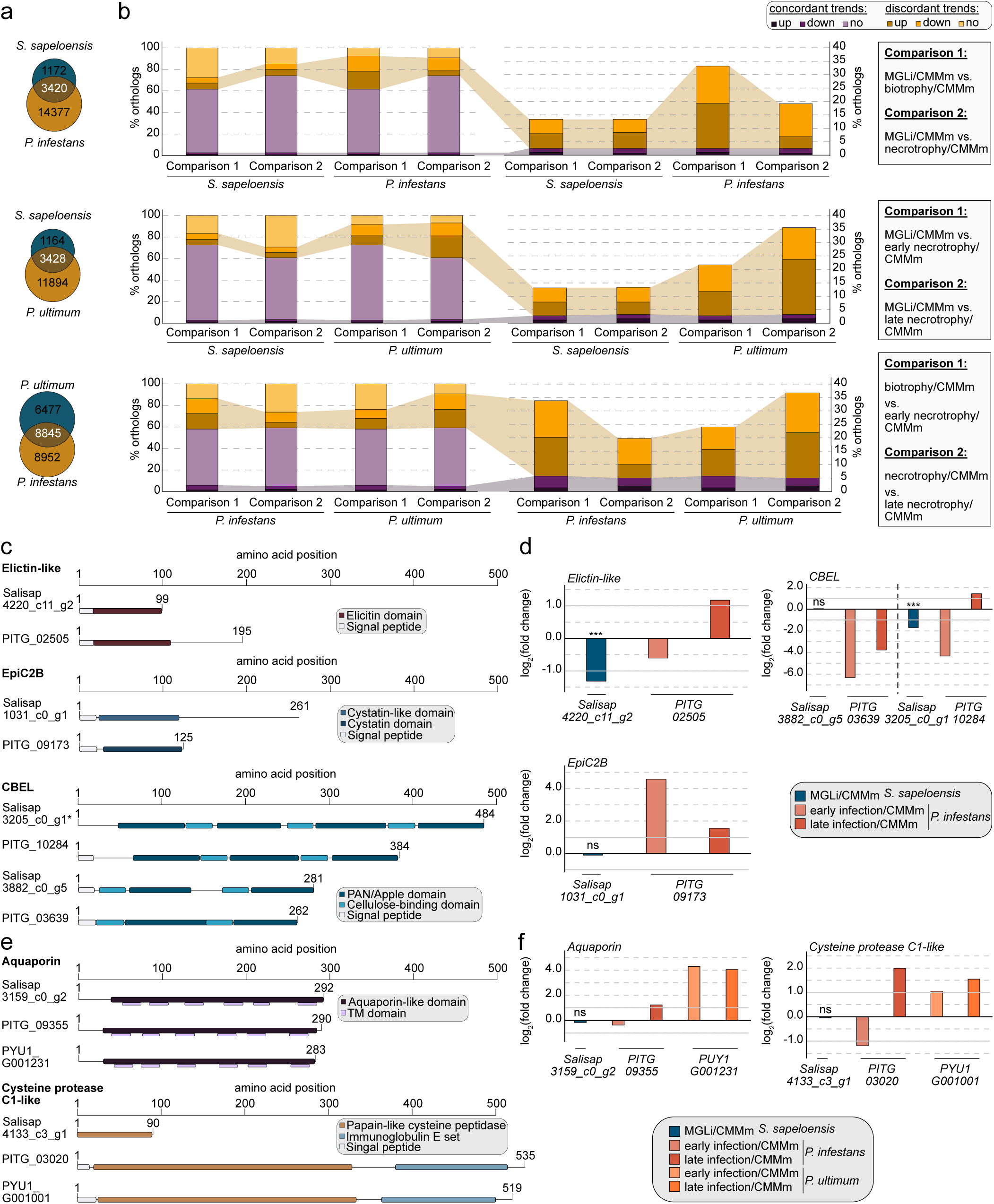
Comparative transcriptomics of *Salisapilia sapeloensis* with two oomycete plant pathogens. (a) The Venn diagrams show the shared orthologs between *P. infestans, P.ultimum* and *S. sapeloensis*. (b) The bar graphs give the percentage concordantly (purple) and discordantly (orange) regulatory trends of orthologs between *S. sapeloensis* and *P. infestans* (top), *S. sapeloensis* and *P. ultimum* (middle) and *P. infestans* and *P. ultimum* (bottom). We compared regulatory trends of orthologs in *S. sapeloensis* MGLi vs. CMMm with early infection vs. CMMm (Comparison 1) or with late infection vs. CMMm (Comparison 2). In the comparisons between the two pathogens Comparison 1 corresponds to early infection vs CMMm of *P. infestans* with early infection vs. CMMm of *P. ultimum*. Comparison 2 corresponds to the comparisons between the late infection phases. Up indicates that a gene is induced (log_2_[FC]≥1), down indicates that a gene is reduced (log_2_[FC]≤−1) and no indicates that a gene shows no changes (−1>log_2_[FC]<1). Right: Overview of all orthologs. Left: Overview of orthologs that are induced or reduced. (c) Domain structure of the *in silico* translated pathogenicity-associated genes encoding Elicitin-like, EpiC2B and CBEL proteins of *S. sapeloensis* and their orthologs in *P. infestans*. The asterisk indicates a partial N-terminus. (d) log_2_(FC) of Elicitin-like-, EpiC2B- and CBEL-encoding genes based on transcriptomic data. The cutoffs for induction and reduction as defined above are indicated with the grey lines at log_2_(FC) = −1 and log_2_(FC) = 1. For *S. sapeloensis* the significant differences are indicated as not significant (ns) when FDR>0.05, and as significant *** FDR≤0.001. The statistical evaluation corresponds well with the previously defined cutoff. (e) Domain structure of *in silico* translated pathogencity-associated genes that show similar regulatory trends in the two pathogens but are differently expressed in *S. sapeloensis*. The structure is given for the orthologs in *S. sapeloensis, P. infestans* and *P. ultimum*. (f) log_2_(FC) of Aquaporin- and Cystein protease-encoding orthologs of *S. sapeloensis, P. infestans* and *P. ultimum*. The cutoffs for induction and reduction of gene expression (log_2_[FC] = − 1 and log_2_[FC] = 1) are indicated as grey lines in the bar diagram. Ns indicates not significant based on the statistical evaluation in *S. sapeloensis*’ transcriptome.

To identify how similar the expression between *S. sapeloensis* and the two pathogens is when comparing data from infection vs. litter colonization, we calculated how many genes showed concordant trends in expression. First, we identified orthologs between *S. sapeloensis, P. infestans* and *P. ultimum* (Figure 5a). Second, we calculated the log_2_(FC) between MGLi and CMMm for *S. sapeloensis* and between infection and CMMm for the two pathogens. Using these data, we asked if the expression of orthologous genes in the three species was higher (up; log_2_(FC) ≥ 1), lower (down; log_2_(FC) ≤ −1) or similar (no change; −1 < log_2_(FC) < 1) upon the encounter of plant tissue compared to CMMm. An ortholog was counted as showing a concordant trend in two species if expression was higher (up) in both species, lower (down) in both species or similar (no change) in both species (Figure 5b). We further analysed concordant trends in expression between litter colonization and an early infection phase and between litter colonization and a late infection phase. This was done to account for the fact that *P. infestans* changes from biotrophy to necrotrophy during infection, which is also accompanied by a switch in expression [23].

Overall, *S. sapeloensis* and the pathogens had similar patterns of gene expression (62.0±5.9%; Figure 5b; Dataset S5a). Hereby, the early necrotrophic phase of the two pathogens, (i.e. in the hemibiotroph *P. infestans* the late infection stage; in the necrotroph *P. ultimum* the early infection stage) showed more concordant trends to saprotrophy than the biotrophic (early infection stage of *P. infestans*) and late necrotrophic (late infection stage of *P. ultimum*) phase (Figure 5b). These similarities are not surprising—yet they are confirmatory of our approach and align with what we know about the biology of the lifestyles: Since in both saprotrophy and necrotrophy the oomycete is eventually confronted with dead tissue, it is only logical that the transcriptional response to these cues is partially overlapping. In contrast the transition between biotrophy to necrotrophy requires a switch from a transcriptional program that is outlined to keep a host alive to a transcriptional program outlined to kill a host. In agreement hemibiotrophic pathogens show strong transcriptional shifts between their biotrophic and necrotrophic phases [23,37,38]. Hence, a strong transcriptional difference between a saprotrophic interaction, in which *S. sapeloensis* feeds on dead plant tissue, compared to the biotrophic phase in the hemibiotroph *P. infestans*, where its physiology is optimized to keep the host alive, is only expected.

The main constituent of the orthologs with concordant regulatory trends between the pathogens and the saprotroph were those that showed no change (Figure 5b). Only 0.69-1.85% of the concordantly regulated orthologs were also responsive (up- or down-regulated) upon colonization or infection. Hence a large fraction of the litter-responsive transcriptome of *S. sapeloensis* has distinct regulatory trends compared to the two pathogens. This means that those genes whose function is associated with the degradation of litter are not induced during infection of a plant pathogen, albeit that both the saprotroph and the pathogen have a functional copy of the respective gene. Conducting the comparison between *P. infestans* and *P. ultimum* we found that the relative amount of concordant regulatory trends between the two pathogens was two times higher than what was observed for the comparison between saprotroph and either pathogen (Figure 5b). This indicates that the signaling required by two different pathogens for the infection of a living plant is more similar to each other than the signaling required for the colonization of dead plant material by a saprotroph.

To gain insight into the responsiveness of genes that show discordant trends, i.e. genes that are uniquely-regulated in the organisms, we first identified the uniquely regulated genes and second asked whether they are induced or reduced or showed no change in expression between infection or colonization vs. CMMm (Dataset S5a-c). Of all orthologs identified in *P. infestans* and *S. sapeloensis* as well as *P. ultimum* and *S. sapeloensis*, 32.7±6.2% were uniquely regulated across all four comparisons. This corresponds to 938±178 genes. The pathogens had more uniquely regulated genes that were induced or reduced compared to the saprotroph: 16.8%±3.6% (152±11 genes) of all uniquely-regulated genes were induced or reduced in the saprotroph colonizing the litter (MGLi) vs. CMMm, while 37.1%±10.2% (354±131 genes) were induced or reduced in the pathogens infecting their hosts vs CMMm (Dataset S5a).

On balance, our data show that *S. sapeloensis* has distinct gene expression compared to the two oomycete pathogens. Such differences in expression suggest that the underlying gene regulatory networks of *S. sapeloensis*, respond differently to the colonization of dead plant material than those of *P. infestans* and *P. ultimum* do towards infection. Next stands the question of which genes are differently regulated in the saprotroph and the two pathogens and whether those genes show a connection to virulence.

### Pathogenicity-associated genes show different expression profiles in *Salisapilia sapeloensis* and two plant pathogenic relatives

To identify genes whose expression differs between *S. sapeloensis* and the pathogens, we investigated the regulatory trends of genes that are shared by *S. sapeloensis* and the pathogens *P. infestans* and *P. ultimum*. To identify unique profiles that set the pathogens apart from the saprotroph, we focused on the top-100 up-regulated orthologous genes with discordant regulatory trends (Dataset S5b). Among these genes, we found an *elicitin-like* candidate (*Salisap4220_c11_g2*). Its putative protein product was slightly shorter than its ortholog in *P. infestans*, but possessed both a signal peptide (SP) and Elicitin domain similar to the ortholog of *P. infestans* (Figure 5c). This *elicitin-like* gene was uniquely induced in the late phase of *P. infestans* (log_2_(FC) > 1), while it was down-regulated in *S. sapeloensis* colonizing litter (log_2_[FC]=-1.3, FDR=3.5*10^−7^, Figure 5d). Likewise, the protease inhibitor *EpiC2B* (*Salisap1031_c0_g1*), a virulence factor of *P. infestans* [39], was uniquely induced during early and late infection but showed no change in expression in *S. sapeloensis* in MGLi compared to CMMm (Figure 5d). Despite the different regulation, structural analyses revealed that the putative protein product of the *EpiC2B* ortholog of *S. sapeloensis* was similar in structure compared to *Pi*EpiC2B. It has a cystatin-like domain similar to the cystatin domain of *Pi*EpiC2B (Figure 5c). Both are located N-terminally right behind the SP and are of similar length. These results show that the saprotroph *S. sapeloensis* generally possesses pathogenicity-associated genes. However, their expression is different from those in pathogens. This raises the question whether *S. sapeloensis* could also act as an unrecognized, possibly latent pathogen, but uses different gene regulatory networks during its saprotrophic interaction.

We investigated further whether *S. sapeloensis* expresses other members of pathogen-associated gene families and whether those showed discordant or concordant regulatory trends. We parsed our dataset of detected orthologs (Dataset S3b) for (i) all protease inhibitor-encoding genes (*Epi* and *EpiC*), (ii) *elicitin-like* sequences, (iii) members of the *Nep1-like* (*NLP*) family, and (iv) Cellulose binding elicitor lectin (CBEL)-encoding genes based on annotations in the *P. infestans* genome. All of these families have been shown to induce disease symptoms and/or are required for successful plant infection [39,40,41,42,43,44,45]. This survey revealed expression of a mere two *CBEL* orthologs (*Salisap3205_c0_g1* and *Salisap3882_c0_g5*) in *S. sapeloensis* (even when considering those transcripts with a TPM<1). We detected no orthologs for *NLPs* or other elicitin-, Epi-, and EpiC-encoding genes.

The putative protein products of the two *SsCBEL* orthologs are similar in structure to their counterparts in *P. infestans* (Figure 5c). Salisap3205_c0_g1 has four PAN/Apple domains distributed across the entire protein sequence. Cellulose-binding domains alternate with the PAN/Apple domains, supporting its putative function as a CBEL protein. It however lacks a SP. This is likely due to the sequence being partial at the N-terminus. The second *SsCBEL* ortholog, *Salisap3882_c0_g5*, translates *in silico* into a protein with two PAN/Apple domains and two Cellulose binding domains, each located in front of a PAN/Apple domain. Additionally, we observe an N-terminal SP.

Like the *Epic2B* and *elicitin-like* orthologs, the two CBEL orthologs showed unique regulatory trends (Figure 5d). One of the orthologs was reduced in *P. infestans* during infection (log_2_[FC] < −1), while *S. sapeloensis* showed no change in expression on litter and CMM. The other CBEL-encoding ortholog was specifically induced in the necrotrophic phase of *P. infestans* (log_2_[FC] > 1), but down-regulated in *S. sapeloensis* (FC_log2_=-1.7, FDR=3.1*10^−17^Figure 5d). These data show that on the one hand *S. sapeloensis* does express, and hence encode, members of gene families associated with a pathogen’s ability to successfully infect a living plant host. On the other hand, only few of those genes are expressed and if so, they show discordant expression patterns in *S. sapeloensis* and *P. infestans*. Overall this allows for the possibility that *S. sapeloensis* is capable of a pathogenic interaction but it is not using these factors during saprotrophy. Alternatively, the protein products of these pathogenicity-associated genes could be co-opted for a different use in *S. sapeloensis*.

### *Salisapilia sapeloensis* expresses few RxLR and SSP effector candidates

To be able to infect living plants, pathogens also counteract the plant immune system by secreting proteins, called cytoplasmic effectors, into the host cells or their apoplast [46,47]. We have already investigated the expression of several of such pathogenicity-associated genes in the previous sections. Here we now focus on Crinklers (CRNs), RxLRs and small secreted peptides (SSPs) that are present in several oomycetes [48,49,50]. Yet, only few orthologous effectors exist between the different oomycetes [20,51]. We therefore used a heuristic and HMM search to identify initial candidates for RxLRs and CRN. For SSPs we used EffectorP2.0 [52]. Because these are secreted, cytoplasmic effectors we further determined candidate genes without a transmembrane domain (TM domain) and with a SP.

Using this approach, we identified four RxLR and 27 SSP and no CRN candidates across the oomycete-affiliated and orphan gene dataset (Figure 6a). One putative RxLR was also annotated as an SSP candidate, resulting in 30 effector candidates. In total, 10/30 effector candidates showed a significant up-regulation (FDR ≤0.05) in association with litter (MGLi vs. MGLw and/or vs. CMMm) (Figure 6b, Dataset S6).

**Figure 6.**
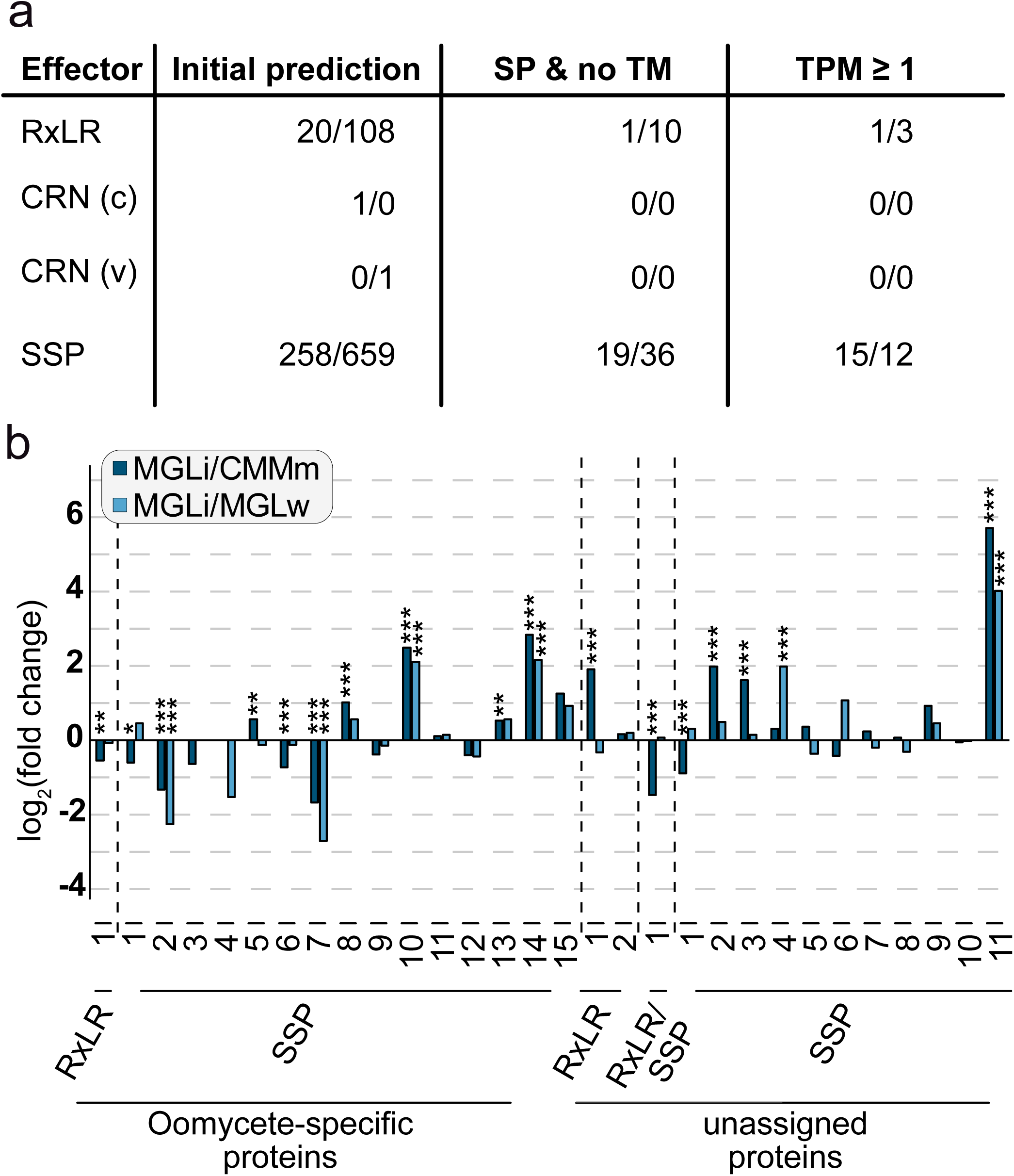
Effector candidate repertoire and expression profile in *Salisapilia sapeloensis*. (a) Number of identified effector candidates in the oomycete-affiliated (left of ‘/’) and orphan gene (right of ‘/’) datasets. On the left the type of effector candidate is given: RxLR, CRN with the canonical CRN motif (c), CRN with an alternative CRN motif (v), and SPP candidates. The table shows the number of effector candidates identified in the ‘initial prediction’ (i.e. compiled results of the HMM and heuristic searches), after filtering for the signal peptides (SP) as well as requiring the absence of transmembrane domains (TM), and after applying an expression cutoff of TPM_TMMnormalize_≥1 in at least one treatment. (b) Relative expression [log_2_(FC)] of effector candidates detected in *S. sapeloensis*. Significant up-/down-regulation is indicated by * FDR ≤ 0.05, ** FDR ≤ 0.01 and *** FDR ≤ 0.001.

Thus, overall, *S. sapeloensis* expressed a few cytoplasmic effector-encoding genes. This further supports the possibility that *S. sapeloensis* could be an opportunistic or latent pathogen. Intriguingly, and in contrast to the above identified pathogenicity-associated genes, some of the cytoplasmic effector-encoding genes were up-regulated in *S. sapeloensis*. This is in agreement with an early up-regulation of some effector-encoding genes in *P. infestans* [35] and might be related to priming for infection of a possible host.

### Common gene expression patterns of Phytophthora infestans and Pythium ultimum differ from those in Salisapilia sapeloensis

To gain more insights into genes that show a saprotroph- or pathogen-associated regulation of gene expression, we honed in on those genes that we identified as having a unique regulation in either *S. sapeloensis* or both pathogens. We asked whether we can identify genes that are induced in both pathogens during infection, but not during colonization of litter; we reason that those genes might have a wider-distributed role in pathogenicity. Here we focused again on the top-100 induced genes in any of the four conditions (*P. infestans* and *P. ultimum*, early and late infection phase; Dataset S5b) and compared them to all uniquely induced genes in the corresponding condition of the other pathogen (i.e. *P. infestans* early infection, vs. *P. ultimum* early infection). We counted only those that are induced in the same condition in the two different pathogens.

Among those genes uniquely induced in either of the conditions we again found nutrient transporter-and CAZyme-encoding genes (Dataset S5b), adding further evidence to a distinct degradation profile of the pathogens and the saprotroph. Apart from the nutrient transporters and CAZymes, we identified aquaporin-encoding genes as well as a cysteine protease family C1-related-encoding gene (Dataset S5b). Aquaporins are also a type of transporters, they allow water permeability across membranes, but are also implicated in transport of for example reactive oxygen species or glycerol [53,54]. All three aquaporin candidates possess an Aquaporin-like domain spanning the entire protein as well as six TM domains (Figure 5e). The latter speak for their integration in the oomycetes’ membranes. Aquaporins are involved in pathogenicity in fungal, apicomplexan and bacterial pathogens [53,54,55]. The induction of the aquaporin-encoding genes (*PITG_09355* and *PYU1_G001231*) in the two distinct oomycete pathogens during infection suggests some importance during the infection process of these oomycetes (Figure 5f). This is corroborated by the contrasting response in *S. sapeloensis* (*Salisap3159_c0_g2*; Figure 5f). Cysteine proteases from hosts and pathogens act in the plant-pathogen interface [56]. In oomycetes C1 family proteases are also present in the *in silico* secretomes of the pathogens [49]. In agreement, the putative protein products of the two pathogens encode a SP (Figure 5e). We could not detect a SP in *S. sapeloensis*’ ortholog. However, we noted that the sequence was rather short, seemed to include only a partial Papain-like cysteine peptidase domain and was missing the Immunoglobulin E set domain (Figure 5e). It is therefore likely that it is a partial sequence. Similar to Aquaporins, an induction in both *P. infestans* and *P. ultimum* during late infection supports a role in pathogenicity (Figure 5f).

The differences in gene expression patterns suggest that *S. sapeloensis* and the pathogens invoke different gene regulatory networks. In support of this we also found a C3H transcription factor (TF) that was specifically induced in both pathogens during an early infection phase vs. CMMm, but its ortholog (*Salisap4203_c0_g1*) was not differentially expressed in *S. sapeloensis* in MGLi vs. CMMm (Dataset1d). TFs are upstream in a hierarchical gene regulatory network. Even slight changes in their expression can result in i) strong changes of in the expression of their target genes or ii) activation of entirely different gene regulatory networks. The latter can be the case if a TF interacts with other TFs, for example by forming a complex with them: Binding specificity or affinity is defined by the combination of TFs [57]. If the expression of the regulatory complex is no longer co-opted, other TF-TF interactions may occur, which in turn will cause a different down-stream expression pattern.

### Gene expression of TF-encoding genes is associated with saprotrophy in *Salisapilia sapeloensis*

*P. infestans* and *P. ultimum* are only two pathogens and more closely related to each other than to *S. sapeloensis*. Additionally, the transcriptomic data that we used to estimate the concordant regulatory trends was generated from tuber infections, not leaf. For a more unbiased view, we hence extended our analysis to eight oomycete pathogens with three different pathogenic lifestyles (four biotrophs, three hemibiotrophs, one necrotroph), five different hosts and four different tissue types (Figure 7; Dataset S5d,e). As discussed above, minor switches in TF binding or expression may alter expression of many genes simultaneously. Because of the distinct expression patterns observed in the previous analyses, we therefore focused on the candidate genes encoding for TFs.

**Figure 7.**
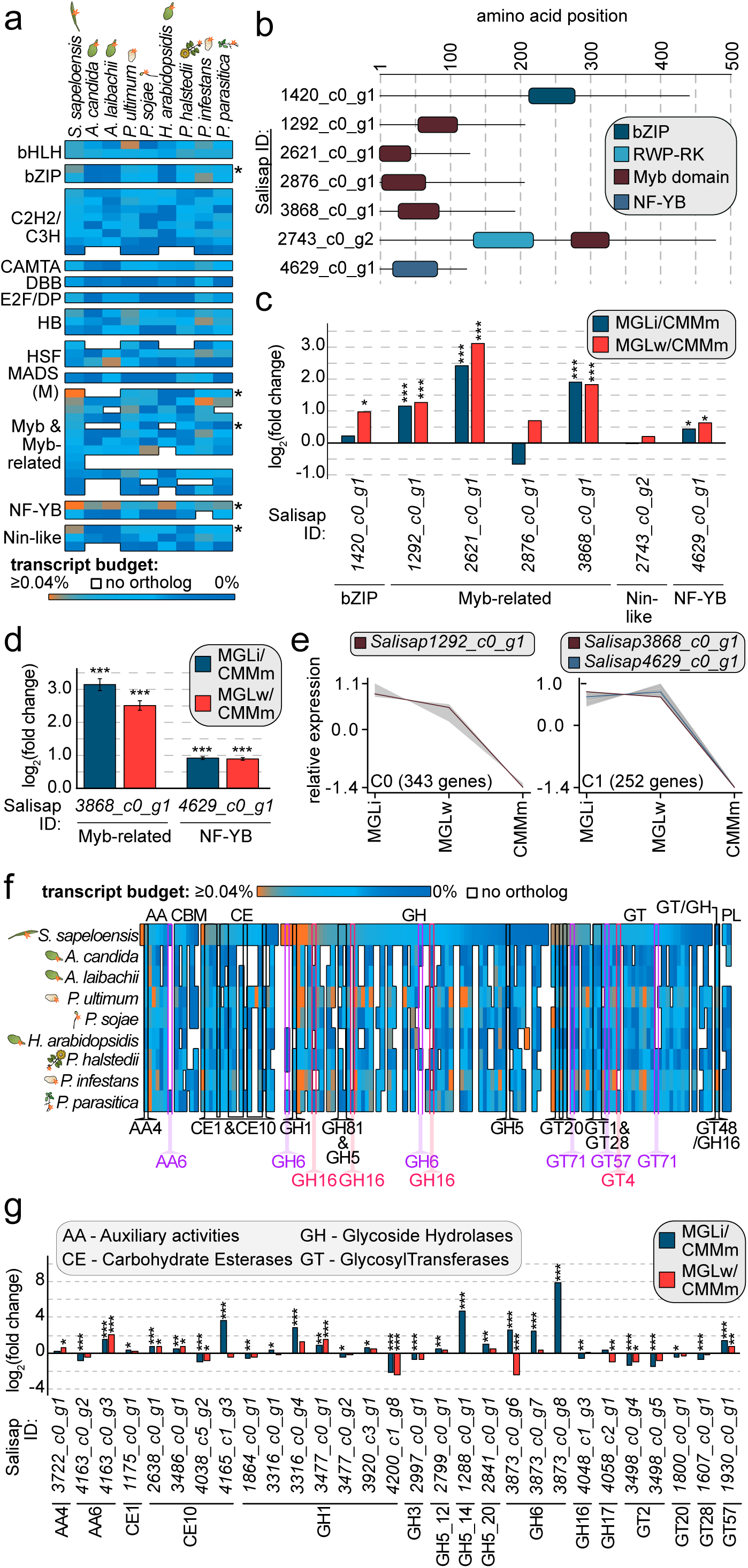
Transcript budget analyses of transcription factor- and CAZyme-encoding genes across plant colonizing oomycetes. (a) Transcript budget (given in %TPM of the total TPM) invested into TF-encoding genes from dark blue (0% transcript investment) to orange (≥0.04% transcript investment). Every line indicates a TF-encoding gene in *S. sapeloensis* and its orthologs in eight plant pathogenic oomycetes. White spaces indicate that no ortholog was identified. The heatmap is sorted according to the transcript budget in *S. sapeloensis* from high to low and per TF category. Asterisks show those TF-encoding gene with an increased transcript budget (log_2_[FC] ≥1) in *S. sapeloensis* vs. all oomycetes with an ortholog. The tissue that was colonized/infected by the oomycete is indicated with the symbol to the right of their names. (b) Domain structure of the *in silico* translated seven TF-encoding genes with increased transcript budget or no orthologs in the plant pathogens. (c) log_2_(FC) of the seven TF-encoding genes in MGLi (blue) or MGLw (red) vs. CMMm in the transcriptome data. Significant differences are indicated as * FDR ≤ 0.05, ** FDR ≤ 0.01, *** FDR ≤ 0.001. (d) qRT-PCR of a Myb-related- and the NF-YB-encoding gene from the seven candidate TF-encoding genes. The average log_2_(FC) for MGLi (blue) or MGLw (red) vs. CMMm is shown as a bar diagram. Error bars indicate the SEM. The reference gene was *SsH2A*. Significant differences are given as *** p-value ≤ 0.001. (e) The clusters C0 and C1 include three of the five TF-encoding genes with an up-regulation in MGLi or MGLw vs. CMMm. Myb-related-encoding genes are indicated with a dark red line and the NF-YB-encoding gene is indicated with a dark blue line. (f) Comparison of the transcript budget invested in CAZyme-encoding genes. The heatmap shows the genes from *S. sapeloensis* and their respective orthologs in eight plant pathogens that infect five different hosts and four different tissues (symbolized by the icons to the left of the species name). Heatmaps are sorted within each CAZyme category and from high transcript investment (orange) to low (dark blue) in *S. sapeloensis*. Each line represents one gene in the saprotroph and its orthologs in the pathogens. White indicates that no ortholog was identfied in a given pathogen. Highlighted CAZyme-encoding genes have a log_2_(FC)≥1 difference in %TPM between either *S. sapeloensis* compared to all pathogens or vice versa. (g) log_2_(FC) of gene expression of all CAZyme-encoding genes with increased transcript budget or no orthologs in other oomycetes. Only genes with a sufficient expression in all conditions (TPM_TMM-normalized_≥1) are shown. Significant differences are indicated as * FDR ≤ 0.05, ** FDR ≤ 0.01, *** FDR ≤ 0.001.

The transcriptome of *S. sapeloensis* featured 47 TF-encoding genes, 40 of which had a TPM_TMM-normalized_ ≥1. These 40 TFs are from 14 TF families. The genomes of the eight pathogen species encoded on average 35±3 TFs that were orthologous to the 40 TFs of *S. sapeloensis*. We detected the most TF orthologs in *P. ultimum* (38) and the least in *Albugo laibachii* (30). Two TF candidate genes of *S. sapeloensis*, both belonging to the Myb-related family, had no ortholog in any of the eight plant pathogens (Figure 7a). Five TF candidate genes were, when its orthologs were present in the pathogens, more highly expressed in *S. sapeloensis* than in the pathogens (log_2_[FC] ≥ 1; Figure 7a; Dataset S5d). These included a bZIP-, two Myb-related-, a NF-YB-, and a Nin-like-candidate. On the flip side, no TF-candidate had a consistently higher expression across all pathogens compared to *S. sapeloensis*.

In view of the large number of genes that are responsive to either colonization or infection and exhibit discordant regulatory trends, the TF data warrants attention. Together, these data suggest that gene regulatory networks are differently wired during saprotrophy and pathogenicity. In this context it can either be that i) similar components of the networks (i.e. if orthologs exist) have to interact differently, because of the distinct expression levels or ii) new components can be added or can replace existing factors in the gene regulatory networks. Both cases will alter downstream expression patterns, such as we have observed, and are not mutually exclusive. Our data on transcript investment in TF-encoding genes across several pathogenic oomycetes and *S. sapeloensis* hence represent additional support for the usage of slightly altered gene regulatory networks by the saprotroph and the pathogens. The data further give us seven TF candidates that we analysed in more depth. Domain predictions using Interpro (Figure 7b) identified one Myb domain in the four Myb-related TF candidates, which is typical for Myb-related TFs [58,59]. Likewise, we confirmed the presence of the bZIP domain in the bZIP candidate, the NF-YB domain in the NF-YB candidate and one RWP-RK and Myb domain in the Nin-like candidate, typical for these TFs [59,60]. Interpro further predicted DNA binding sites in all seven candidates.

Higher transcript investment does not necessarily mean that those genes are also up-regulated during litter colonization, therefore we investigated their expression in the *S. sapeloensis* transcriptome. Of the seven TFs, five were up-regulated in either MGLi or MGLw vs. CMMm (FDR ≤0.05). The other two showed no differential expression in these comparisons. Of the five TF-encoding genes, the bZIP-encoding gene was only up-regulated in MGLw vs. CMMm. while three Myb-related-encoding and the NF-YB-encoding gene were up-regulated in both MGLi and MGLw vs. CMMm (Figure 7c). This also included the two Myb-related-encoding genes that had no orthologs in the other plant pathogens (*Salisap1292_c0_g1* and *Salisap2621_c0_g1*). We further tested two TF-encoding genes (one Myb-related-encoding gene, *Salisap3868_c0_g1* and the NF-YB-encoding gene *Salisap4629_c0_g1*) using qRT-PCR (Figure 7d). Both genes were up-regulated in MGLi and MGLw compared to CMMm (p-value<0.001). Hence, the qRT-PCR confirmed the transcriptomic data in both cases. Overall, these data suggest that the majority of the TFs we identified with our screening are also important in litter colonization and/or degradation.

To get a better picture of the putative role of our candidate TFs, we used the bioinformatic tool *clust* to predict co-expression patterns of the five TF candidates up-regulated in both MGLi and MGLw vs. CMMm with other genes in our transcriptome. Co-expression analyses are a powerful tool to identify genes involved in the same biological processes, including the potential regulatory relationships between TFs and candidates for downstream target genes [61,62,63,64,65]. In most cases, co-expression analyses consider positively co-expressed genes. The caveat of such analyses with regard to TF-target relationships is that it can only identify TFs that are positive regulators with immediate impact on the expression of the target genes. The idea is that in these cases the target gene has a similar gene expression pattern as it’s regulating TF. Despite this caveat, co-expression analyses have successfully identified regulatory relationships between TFs and target genes [63,65]. As we are currently missing genome data for *S. sapeloensis* we used this approach to get a rough idea of possible target candidates.

In total, the transcriptomic data was broken into 19 clusters of co-expressed genes (C0-C18). These 19 clusters contained 2165 of the 4592 oomycete-affiliated genes (Figure S3); another 1434 genes were not co-expressed with a sufficient number of genes and were hence not sorted into any cluster. Of the five TF-encoding genes, three were found to be co-expressed with other genes in the transcriptome, and two had a unique expression profile. The three genes included two Myb-related (*Salisap1292_c0_g1* and *Salisap3868_c0_g1*) and the NF-YB (*Salisap4629_c0_g1*) candidate genes (Figure 7e).

The Myb-related-encoding gene *Salisap1292_c0_g1* clustered with 342 other genes in cluster C0, while the Myb-related-encoding gene *Salisap3868_c0_g1* and the NF-YB-encoding gene *Salisap4629_c0_g1* are co-expressed with 250 other genes in cluster C1 (Figure 7e). Both clusters encompass genes that exhibit an elevated expression in MGLi and MGLw compared to CMMm. This is in agreement with the up-regulation of the three TF-encoding genes in MGLi and MGLw vs. CMMm (Figure 7c,d). Neither C0 nor C1 showed any GO-term enrichment. We visualized the annotation of all differentially expressed genes that are up-regulated in MGLi vs. CMMm (FDR ≤0.05) for C0 and C1 (excluding all annotations as hypothetical and uncharacterized proteins; Figure S4). Several distinct functional groups were represented in C0 and C1, which explains the absence of any enrichment of ontologies. However, we again noticed the presence of many CAZyme- and transporter-encoding genes among the C0 and C1 cluster. Hence these genes show a similar expression with the investigated candidate TF-encoding genes. Our previous analyses revealed distinct gene expression patterns between *S. sapeloensis* and two plant pathogens, suggesting that degradation and nutrition uptake has pathogen- and saprotroph-specific patterns. This now allows for a correlation of saprotroph-specific expression of TF-encoding genes with the expression of degradation- and nutrient uptake-associated genes.

### A putative link between expression of CAZyme-encoding genes and transcription factor-encoding genes highlighted by comparative transcriptomics

The results from the clustering primed us to further investigate the distribution of CAZyme-encoding genes across the 19 clusters. In total, 72 of the 149 CAZyme-encoding genes are present in a cluster. Of those 20 (27.8%) are found in either C0 or C1 and are hence associated with both litter and litter-associated growth of *S. sapeloensis*. In both clusters are CAZymes from the families AA, CBM, CE, GH and GT, but not PL. In agreement with their clustering most of the genes in C0 (six of seven) and C1 (seven of 13) were differentially up-regulated in MGLi and/or MGLw vs. CMMm (FDR ≤0.05).

We next asked whether CAZyme-encoding genes show a lifestyle-associated transcript investment, as observed for the TF-encoding genes (Figure 7f). To answer this question, we first searched for orthologs of the 149 CAZyme-encoding genes from *S. sapeloensis* in the eight genomes of the plant pathogens. For 24 (i.e. 16.1%) we found no orthologs across the targeted plant pathogens, which suggests that they are either undetectable by this approach due to an extremely high divergence or lineage-specific acquisitions in *S. sapeloensis*. These likely lineage-specific genes were distributed across all CAZyme-categories (except GH/GT and PL). In agreement with the GO-term analyses for genes unique to *S. sapeloensis* (Figure 3a), the majority of them (16 of 24 genes) encoded GHs (Figure 7f, Dataset S5e); these data, again, highlight the unique CAZyme-repertoire of *S. sapeloensis*. In five of the seven categories (AA, CE, GH, GT and GH/GT) we found CAZyme-encoding genes in which either *S. sapeloensis* or all pathogens invest a much higher fraction of the transcript budget (log2 [FC]≥1; Figure 7f, Dataset S5e). This included 26 genes (belonging to AA, CE, GH, GT) with a higher transcript investment in *S. sapeloensis* and one (belonging to GH/GT) with a higher transcript investment in all pathogens. Of the 50 genes that either had higher expression levels in *S. sapeloensis* or had no orthologs in any of the pathogens, 18 were up-regulated in MGLi and/or MGLw vs. CMMm (FDR<0.05, Figure 7g). Taken together, these 18 genes are likely relevant for saprotrophy in *S. sapeloensis* and are used differently in oomycete pathogens.

To link the expression of CAZyme-encoding genes and TF-based gene regulation, we compared the transcript investment data from CAZyme- and TF-encoding genes with the clustering analyses. This pinpointed five CAZyme-encoding genes i) into which *S. sapeloensis* invested much higher transcript levels compared to the pathogens and ii) that were co-expressed with one of the Myb-related- or NF-YB-encoding genes. These five genes include one AA6-, two CE1- and two CE10-encoding genes (*AA6*: *Salisap4163_c0_g3; CE1*: *Salisap3051_c0_g2, Salisap3972_c0_g1*; *CE10*: *Salisap2638_c0_g1*, *Salisap3486_c0_g1*). Each cluster (C0 and C1) contains one CE1 and one CE10 genes. The AA6-encoding gene is clustered in C1. Three genes were sufficiently expressed in all three conditions so that a log_2_(FC) could be calculated (*Salisap4163_c0_g3* [*AA6*]*, Salisap2638_c0_g1* [*CE10*] and *Salisap3486_c0_g1* [*CE10*]). Indeed, all three genes were significantly up-regulated in MGLi and MGLw vs. CMMm (FDR≤0.05; Figure 7g). Overall, this makes the AA6- and the four CE-encoding gene candidates relevant for tissue degradation in oomycete saprotrophy compared to pathogenicity. Additionally, the involvement of the three TFs in regulating the expression of the AA6-encoding and four CE-encoding genes is possible.

### Conclusion

Oomycetes are an exceptionally diverse group of microorganisms. They include many lineages with pathogenic and non-pathogenic lifestyles; lifestyles that evolved multiple times throughout the evolutionary history of the group. This makes oomycetes an excellent group to study the evolution of microbial lifestyles. Comparisons between saprotrophic and pathogenic interactions can reveal vital clues about what, at the molecular and biochemical level, distinguishes a pathogen from a non-pathogen. We have established a first reference point for the differential regulation of gene expression that occurs during oomycete saprotrophy. Investigation of *S. sapeloensis* allowed us to contrast pathogenic and non-pathogenic gene expression patterns. Due to the distinct expression responses to infection and litter colonization we hypothesize that *S. sapeloensis* may have evolved slight alterations in gene regulatory networks. Whether *S. sapeloensis* is capable of invoking gene regulatory patterns associated with pathogenicity remains to be determined. Because the organism possesses pathogenicity-associated factors, we cannot rule out the possibility that it is a latent or opportunistic pathogen (as is also observed for *Aphanomyces stellatus*, for which a genome was also published very recently [50]). Given their environment/host, saprotrophs may be capable of modulating their gene regulatory networks towards either remaining saprotrophic or becoming virulent. We have shown how the evolutionary history of these lifestyles can be studied from the perspective of global gene expression patterns. Such analyses can, in turn, inform the study of extant pathogens by revealing new candidate genes for genetic investigations of lifestyle specificity and change.

## Methods

### Growth conditions and inoculation of marsh grass litter

*Salisapilia sapeloensis* (CBS 127946) was grown on corn meal agar (1.8% sea salt; corn meal extract [60g/l]; 1.5% agar) at 21°C in the dark. *Spartina alterniflora* was collected at 44°40’ N, 63°25’ W, washed, cut in small pieces and autoclaved (marsh grass litter). Autoclaved litter was transferred to 24-well plates containing sterile salt water (1.8% sea salt) and inoculated with agar plugs with 2-6 days old mycelium of *S. sapeloensis*. As a control mycelium of *S. sapeloensis* was transferred to 24-well plates containing liquid CMM (1.8% sea salt). Inoculation and control were incubated in the dark at 21°C for 7days. For each treatment three biological replicates (i.e. three distinct 24-well-plates per treatment) were prepared.

### Staining and Microscopy

Light microscopy was done using an Axioplan II (Axiocam HRC colour camera; Zeiss). Trypan blue staining was performed according to [66]. For laser scanning confocal microscopy (LSM 710 [Zeiss]), mycelium was stained with 1% calcofluor white, prepared from powder (Fluorescent Brightener 28, Sigma Aldrich) solved in water according to [67].

### DNA extraction and identification of marsh grass

DNA was extracted with Edwards buffer [68]. We amplified (PrimeSTAR® Max DNA Polymerase; TaKaRa) the *ribulose-1,5-bisphosphate carboxylase/oxygenase large subunit* (*RbcL*) and the *partial18S-ITS1 (internal transcribed spacer 1)-5.8S-ITS2-partial28S* region using rbcLa-F [69]/ rbcLa-R [70] and rbcLa-F/rbcLajf634R [71] for *RbcL* and ITS2-S2F [72]/ITS4 [73] for the *partial18S-ITS1-5.8S-ITS2-partial28S* region. PCR products were purified (Monarch® PCR & DNA Cleanup Kit 5µg; New England BioLabs) and sequenced from both sides with Eurofins Genomics. After quality assessment, we generated consensus sequences (accessions: MH926040, MH931373). Blastn [74] was used against NCBI nt. The *partial18S-ITS1-5.8S-ITS2-partial28S* sequence retrieved two equally good hits (100% coverage, 99.718% identity, e-value 0): both hits were to *Sporobolus alterniflorus* (synonyme: *Spartina alterniflora*). The *RbcL* sequence retrieved *Sporobolus maritimus* and *Spartina alterniflora* (99% coverage, 100% identity, e-value 0). Of the two only *S. alterniflora’s* range includes Nova Scotia.

### RNA extraction and Sequencing

Biological triplicates per treatment (3x *S. sapeloensis* grown in liquid CMM, 3xsalt water and 3xMGL) were harvested for RNA extraction. For mycelium grown directly in liquid CMM or salt water close to MGL four wells per biological replicate were pooled and ground in 1ml TRIzol^TM^ using a Tenbroeck homogenizer. For *S. sapeloensis* colonizing MGL, 12 wells of litter were combined per one biological replicate and ground in liquid nitrogen using mortar and pestle and 100mg powder was extracted using TRIzol^TM^. RNA was treated with DNAse I. RNA concentration was assessed using an Epoch spectrophotometer (BioTek) and quality was assessed using a formamid gel. The RNA was sent to Genome Québec (Montréal, Québec, Canada), who analyzed RNA using a Bioanalyzer. Nine libraries (biological triplicates, three treatments) were prepared with high-quality RNA and sequenced using the Illumina HiSeq4000 platform for 100 paired-end (PE) reads. In total, we obtained 344,505,280 million paired reads; 312,999,896 million paired reads remained after trimming and filtering for quality (Figure S2).

### Data processing and taxonomic annotation

Quality of the raw data was analyzed using FastQC v. 0.11.5 (www.bioinformatics.babraham.ac.uk/projects/fastqc). Trimming and adapter removal was done using Trimmomatic v. 0.36 [75] (settings: ILLUMINACLIP:<custom_adapter_file>:2:30:10:2:TRUE HEADCROP:10 TRAILING:3 SLIDINGWINDOW:4:20 MINLEN:36). Trimmed reads were re-analyzed with FastQC v. 0.11.5 and are available within Bioproject PRJNA487262. We pooled all read data for a *de novo* assembly using Trinity v. 2.5.0 [76] with Bowtie2 v. 2.3.3.1 [77], producing 44669 isoforms that were taxonomically annotated using DIAMOND blastx v0.8.34.96 [78]: All isoforms were queried against the nr database (June 2017). Hits that were recognized as statistically significant (e-value cutoff 10^−5^) were kept and sorted according to taxonomy. Isoforms with a hit to an oomycete protein were assigned to the oomycete-affiliated dataset.

### Differential expression analyses

Abundance estimates for all different isoforms were calculated using Bowtie2 v. 2.3.3.1 and RSEM v. 1.2.18 [78] and TPM_TMM-normalized_ values were calculated. Differential gene expression was analyzed based on the read counts using edgeR v. 3.20.9 (limma v. 3.34.9; [80]), which models the data as negative binomial distributed and uses an analogous test to the Fisher’s exact test to calculate statistical significance. Afterwards a Benjamini–Hochberg false discovery rate correction was applied. Data from all conditions were hierarchically clustered according to their similarity in per-gene expression change (log_2_-transformed) from the median-centered expression per transcript. A representative isoform (the isoform with the highest expression) was identified, retrieving 8023 unique genes. After a NCBI contamination screening, 7777 genes were deposited at GenBank (Figure S2; Bioproject PRJNA487262).

### *In silico* protein prediction

We predicted protein sequences for all major isoforms i) with a taxonomic affiliation to oomycetes (oomycete-affiliated dataset) and ii) with no hit in the databases (i.e. potential orphan genes [unassigned dataset]). For the oomycete-affiliated dataset, we predicted all possible reading frames using EMBOSS’ getorf [81] and then used blastx [74] against i) a database containing all hits retrieved from the DIAMOND blast output and ii) an oomycete proteome database consisting of the protein sequences of 22 oomycete genomes (Table S1) using the unique genes as query. Based on the best blast hit we determined the most-likely open reading frame and protein sequence. For the unassigned dataset, we used the bioperl script longorf (https://github.com/bioperl/bioperl-live/blob/master/examples/longorf.pl) with the option *strict*.

### Functional annotation

For functional annotations we used EggNOG-mapper 4.5 [82] and GhostKOALA [83], with the KEGG database *genus prokaryotes and family eukaryotes*. Further, we performed a blastp [74] search against our DIAMOND output and oomycete proteome database (e-value cutoff of 10^−5^). Differentially expressed genes were manually annotated using these data. If databases produced contradicting results, we used CD search [84] to identify protein domains within the predicted sequences and chose the better supported annotation.

### Prediction of putative CAZymes

To predict the CAZymes of *S. sapeloensis* and 22 oomycetes we used HMMER3 [85] on dbCAN [86]. Sometimes the same sequence was predicted to belong to more than one family of CAZymes. We divided those cases in two groups: Group I contains sequences with HMM-hits with different functional specificity (e.g. CBM and GH domains). Here we retained the more specific annotation (i.e. GH, not CBM). Group II contains sequences that had hits with similar functional specificity (e.g. GT and GH). Here we used CD search to validate the presence of the domains [84]. If only one of the two domains was found using this approach, we changed the annotation accordingly. Otherwise, we kept both annotations (category *mixed*).

### CAZyme family enrichment

To analyze the distribution of CAZyme families, we calculated the percentage of CAZymes in the transcriptome of *S. sapeloensis* and 22 oomycetes genomes ([13,20,87,88,89,90,91,92,93,94,95]; Table S1) with different lifestyles and hosts. We compared the CAZyme profile of *S. sapeloensis* with that of the other oomycetes’ genomes. We defined enrichment of expression of CAZyme subfamilies as log_2_(CAZyme-subfamily*_Salisapilia_*[%] / (Ø CAZyme-subfamily_other oomycetes_+SD) [%]) ≥1.

### Prediction of transcription factors

Transcription factors (TF) were predicted using PlantTFDB 4.0 [96] with the oomycete-affiliated dataset as input. This database uses HMMER-based approach to identify TF domains based on Pfam and published domains to identify TFs and can therefore be used outside of the plant kingdom to identify common eukaryotic TFs.

### Prediction of putative effector proteins

Putative effectors were predicted using the oomycete-affiliated and the unassigned dataset, since effectors often show low conservation between species [20] and can be orphan genes. To predict RxLR-dEER type effectors we used an HMMER-based search using HMMER 3.1b1 [85] with the HMM-profile from [51] and a heuristic search identifying the RxLR-dEER motif in the first 150 amino acids [46]. The putative Crinkler type effectors were identified as in [49] using a heuristic search for LFLA[R/K]X and LYLA[R/K]X between the residues 30 to 70. SSPs were predicted using EffectorP 2.0 [52]. Only considering sequences with a TPM_TMM-normalized_ >1, we further required the absence of TM domains (i.e. no domain predicted by either TMHMM v. 2.0 [97], Phobius [98] or TOPCONS2 [99]) and the presence of a SP (predicted by two algorithms independently).

To predict the SPs, we used SignalP3-HMM [100], SignalP4.1-HMM [101], Phobius [98] and TOPCONS2 [99]. We used both SignalP3 and SignalP4.1, because SignalP3-HMM can be a better predictor of SPs for oomycete effector proteins [102]. The prediction of SPs requires intact N-termini. We screened for sequences with intact N-termini using a perl script, resulting in 3242 oomycete-affiliated usable protein sequences. For the unassigned protein sequence dataset, an intact N-terminus was a requirement for translation.

### Comparative transcriptomics

Sequence orthologs between *S. sapeloensis* and the eight oomycetes [21,23,95,103,104] (Table S1) were estimated using a reciprocal blastp approach. Protein sequences that found each other reciprocally were designated as orthologs (e-value cutoff = 10^−5^).

We analysed co-regulation of *S. sapeloensis* and other oomycetes during growth on their plant substrate by comparing differential RNAseq data from orthologous sequences of *S. sapeloensis* and the pathogens *P. infestans* and *P. ultimum*. Expression data from the colonization of litter versus growth in liquid CMM was compared to expression data from early and late infection stages in potato tubers versus early and late growth on CMM. The latter was obtained from [23] (Table S1). We asked whether an ortholog shows co-regulation (i.e. both are up-regulated [FC≥1], down-regulated [FC≤−1] or show no change [FC>−1 and <1] in the plant-associated stage compared to the medium growth stage). For comparison, we also asked how much co-regulation the pathogens show to each other during infection versus growth on medium.

We compared RNAseq data between *S. sapeloensis* and eight plant pathogens from the Peronosporales and Albugonales (different lifestyles, different hosts and tissues; Table S1). Some of these oomycetes are biotrophs and cannot grow outside their hosts. Therefore, we calculated the transcript budget (%TPM or %CPM) to estimate how much transcript an oomycete invests into a gene during infection/colonization and compared it between orthologous genes from the different oomycetes.

For *S. sapeloensis* we used TPM_TMM-normalized_ values from litter-colonizing samples. For *P. infestans* and *P. ultimum* we used CPM_TMM-normalized_ values [23] for the late infection stage. For *Albugo candida, Albugo laibachii, Hyaloperonospora arabidopsidis, Phytophthora parasitica, Phytophthora sojae* and *Plasmopara halstedii*, we downloaded transcriptome data for progressed infection stages (Table S1). Reads were analyzed with FastQC v. 0.11.5, trimmed with Trimmomatic v. 0.36 [75] (settings: HEADCROP:10 TRAILING:3 SLIDINGWINDOW:4:20 MINLEN:36) and mapped to the corresponding species’ transcript sequences using RSEM v. 1.2.18 [79] with Bowtie v. 2.3.3.1 [77]. The data from the different oomycete species is made comparable by only using either log_2_(FC) or transcript budget (relative transcript investment). Both of these serve to normalize the data within a given dataset before comparisons across species.

### Protein sequence and structure analyses

Protein domain structures were predicted using CD search [84] and InterPro [105].

### Co-expression analyses

We used clust [106] to study co-expression in our transcriptomic dataset. As input, we used TPM_TMM-normalized_ values and normalized accordingly from all treatments for all 4592 oomycete-affiliated genes in the oomycete-affiliated dataset. The settings were: Cluster tightness 7, minimum cluster size 11, lowly expressed and flat genes were filtered out prior to clustering, using a 25-percentile cutoff. Retained genes were expressed higher than the cutoff in at least one condition.

### qRT-PCR

cDNA was created from 1000ng of RNA from MGLi, MGLw and CMMm using the iScript™ cDNA Synthesis Kit (Bio-Rad). For all treatments we used three biological replicates. We designed primers for the qRT-PCR using NCBI primer BLAST (Table S2). Primers were tested on cDNA from *S. sapeloensis* using the PrimeSTAR® Max DNA Polymerase (TaKaRa). PCR products were sequenced at GENEWIZ (NJ, USA).

The qRT-PCR was conducted on a CFX Connect^TM^ Real-Time System (BioRad). We tested the expression of four CAZyme-encoding genes (*Salisap1930_c0_g1, Salisap3316_c0_g4, Salisap3873_c0_g7* and *Salisap4165_c1_g3*) and two TF-encoding genes (*Salisap3868_c0_g1* and *Salisap4629_c0_g1*) across RNA from all three treatments (MGLi, MGLw and CMMm) with three biological replicates per treatment and three technical replicates per biological replicate. As reference genes we used *H2A* (*Salisap2927_c0_g1*). *H2A* was selected as a reference gene based on its constant expression across all samples in the transcriptome. Its constant expression was further confirmed in the qRT-PCR. A standard curve with the dilutions 1, 1:10 and 1:100 was performed using pooled cDNA from all conditions. We calculated the relative expression of the genes in MGLi and MGLw against CMMm using [107]. To calculate significant differences between expression we first calculated whether the data was normally distributed using a Shapiro-Wilk test [108]. Second, we tested whether the data we compared had the same variation and third, depending on the results we either performed a two-sampled t-test, a Welch two-sampled t-test or a Mann-Whitney U test [109]. All statistical analyses were performed in R v. 3.2.1.

## Acknowledgement

We thank Emma Blanche for technical assistance.

## Author Contributions

Conceptualization, S.d.V.; Methodology, S.d.V.; Validation, S.d.V.; Formal Analysis, S.d.V., J.d.V.; Investigation, S.d.V., J.d.V.; Data Curation, S.d.V., J.d.V.; Writing – Original Draft, S.d.V.; Writing – Review & Editing, S.d.V., J.d.V., J.M.A., and C.H.S.; Visualization, S.d.V., J.d.V.; Funding Acquisition, S.d.V., C.H.S.; Resources, J.M.A., C.H.S.; Supervision, J.M.A. and C.H.S.

## Supplemental Data

**Dataset S1.** Summary statistics for and information on the *de novo* assembly.

**Dataset S2.** KEGG pathway and BRITE hierarchy annotation and responsiveness towards the three different growth conditions of *Salisapilia sapeloensis*.

**Dataset S3.** Pairwise reciprocal BLASTp analyses of *Salisapilia sapeloensis* versus other oomycetes.

**Dataset S4.** CAZyme annotation of the *de novo* assembled transcriptome of *Salisapilia sapeloensis* and the genomes of 22 other oomycetes; expression analysis of CAZyme-encoding genes in *Salisapilia sapeloensis*.

**Dataset S5.** Comparative transcriptomic analyses across diverse oomycete species.

**Dataset S6.** *In silico* translation of the *de novo* assembled transcriptome of *Salisapilia sapeloensis* and expression analysis of putative RxLRs, CRNs, and SSPs of *Salisapilia sapeloensis*.

**Table S1.** Transcriptomic and genomic data used in this study for comparative analyses.

**Tables S2.** Primers used in the qRT-PCRs.

**Figure S1.** Confocal micrographs of *Salisapilia sapeloensis* growing on and in marsh grass litter.

**Figure S2.** Workflow for transcriptome *de novo* assembly and gene identification.

**Figure S3.** *Clust* analysis of the *Salisapilia sapeloensis* transcriptomic dataset across all conditions.

**Figure S4.** Annotation of litter-associated genes in clusters C0 and C1.

