## Supplementary material for "Comparative analyses of saprotrophy in *Salisapilia sapeloensis* and diverse plant pathogenic oomycetes reveal lifestyle-specific gene expression": Figure S1

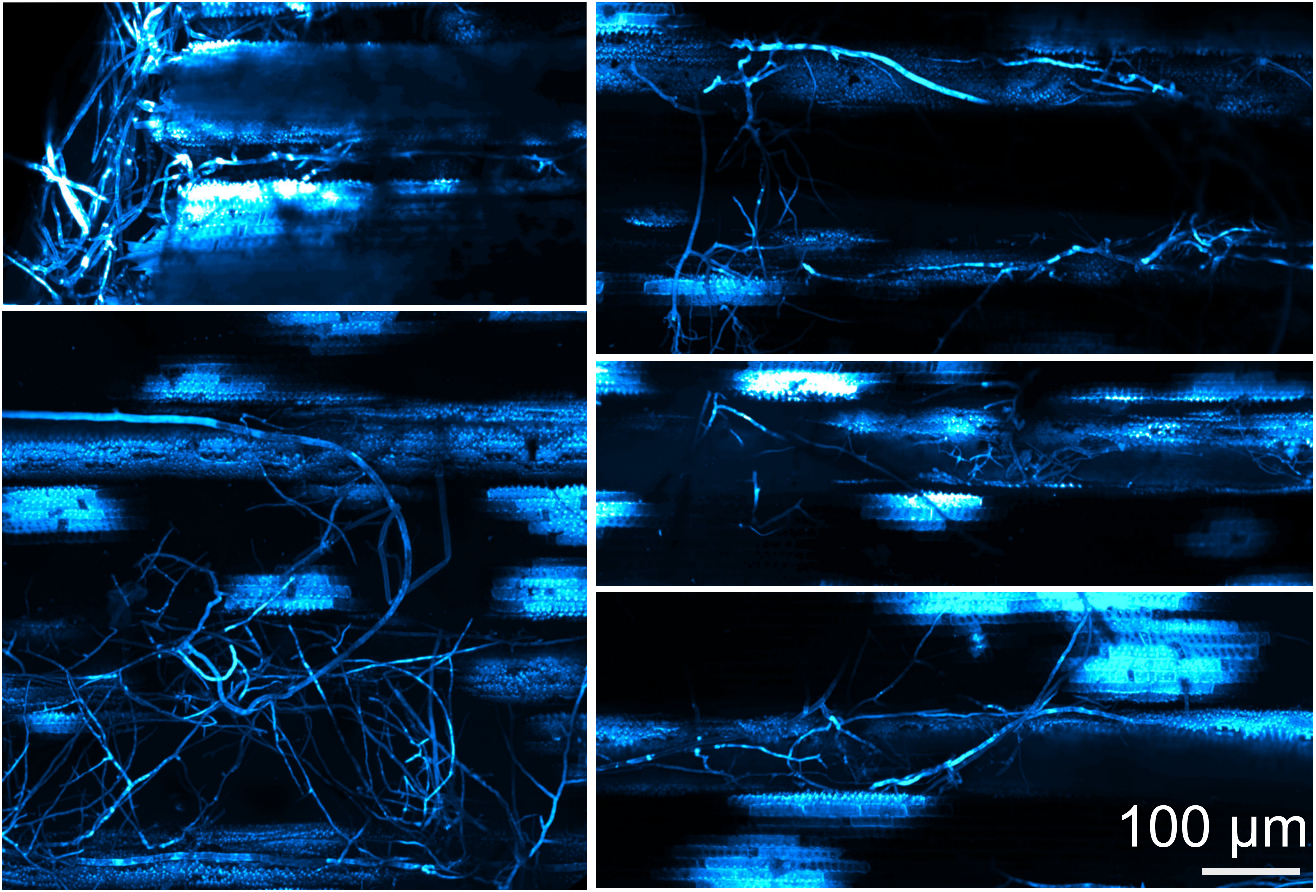


**Figure S1. Confocal micrographs of *Salisapilia sapeloensis* growing on and in marsh grass litter.** Samples were stained with Calcofluor white. Fluorescence is false-coloured in cyan. The z-stacks were reconstructed and projected from whole-mount micrographs. The scale bar applies to all micrographs.
