## Supplementary material for "Comparative analyses of saprotrophy in *Salisapilia sapeloensis* and diverse plant pathogenic oomycetes reveal lifestyle-specific gene expression": Figure S2

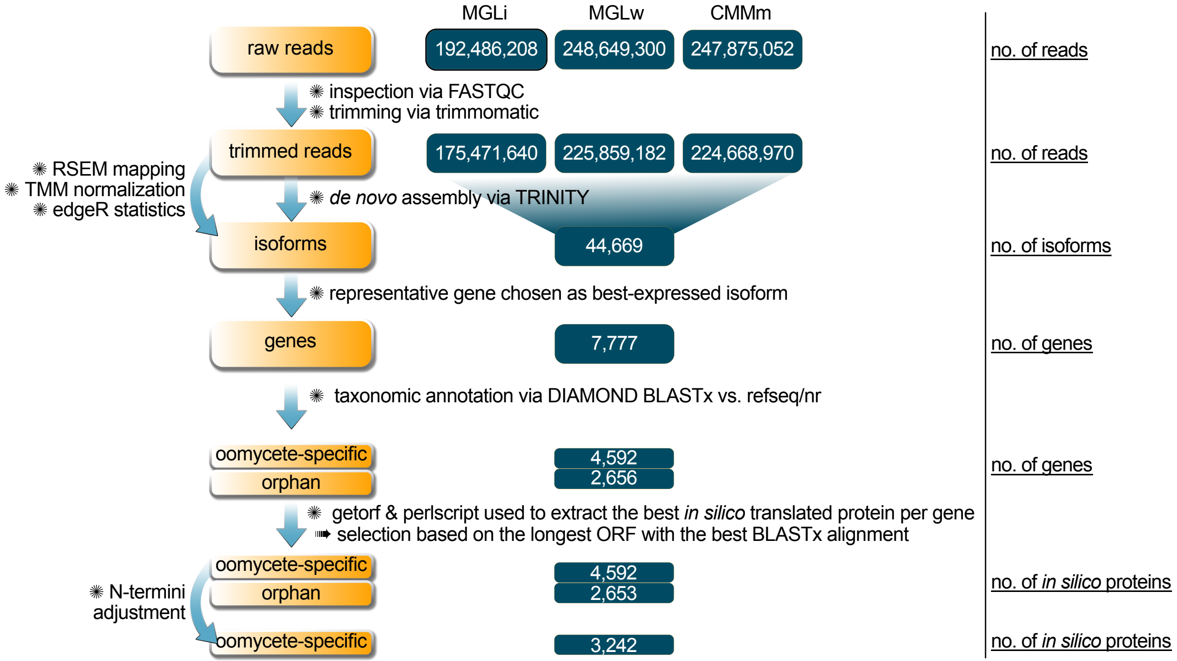


**Figure S2. Workflow for transcriptome *de novo* assembly and gene identification.** MGLi indicates the data from *Salisapilia sapeloensis* grown in litter, MGLw the data from *S. sapeloensis* grown in the water next to litter and CMMm the data from *S. sapeloensis* grown on liquid cornmeal medium.
