## Supplementary material for "Comparative analyses of saprotrophy in *Salisapilia sapeloensis* and diverse plant pathogenic oomycetes reveal lifestyle-specific gene expression": Figure S3

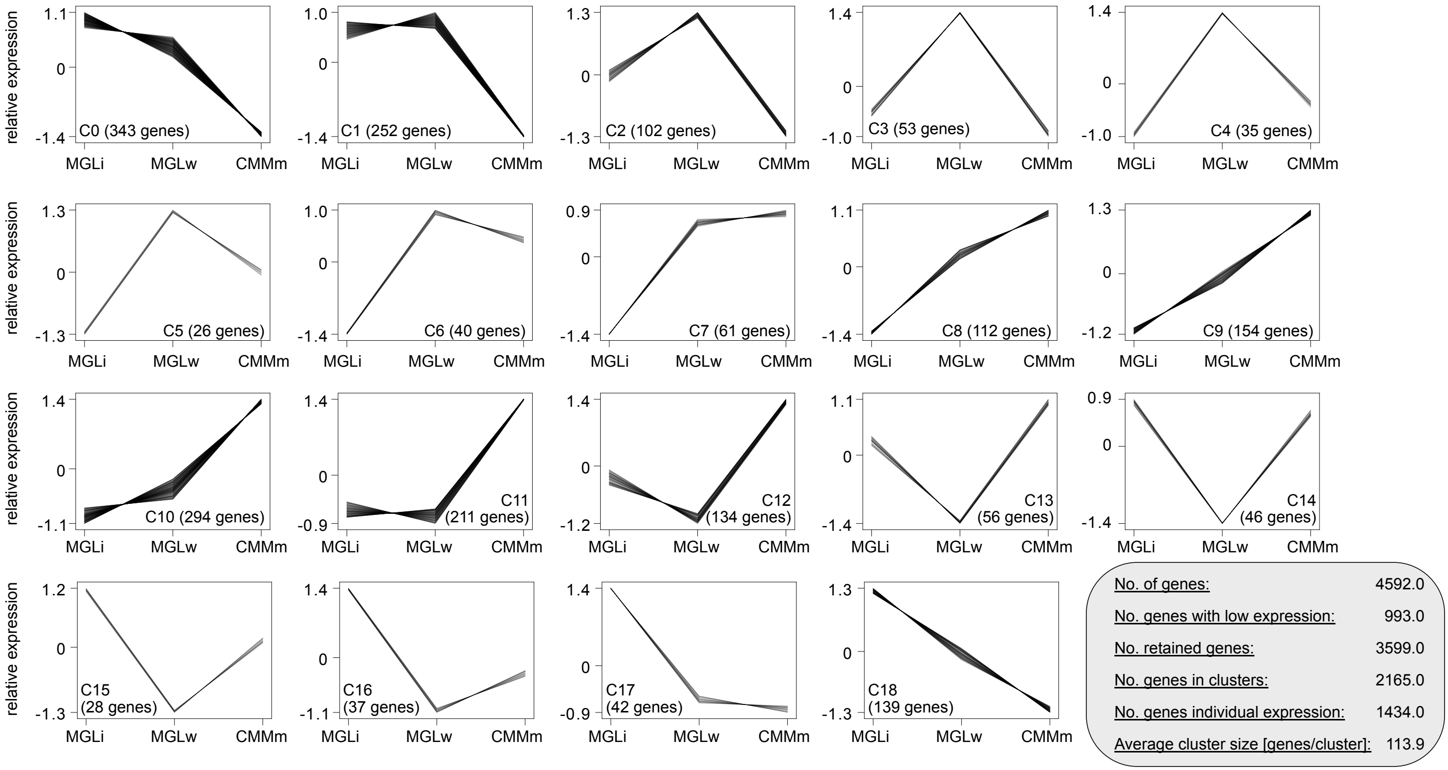


**Figure S3. *Clust* analysis of the *Salisapilia sapeloensis* transcriptomic dataset across all conditions.** The analysis resulted in 19 clusters and includes 2165 of the 4592 oomycete-affiliated genes. The y-axis shows the relative expression based on the $TPM_{TMM\text{-}normalized}$ values and was normalized using a normalization step built-in in the clust analysis. The three conditions are shown on the x-axis. High values indicate a high $TPM_{TMM\text{-}normalized}$ and low values indicate a low $TPM_{TMM\text{-}normalized}$. Genes with low expression in all three conditions were removed from the analysis.
