## Supplementary material for "Comparative analyses of saprotrophy in *Salisapilia sapeloensis* and diverse plant pathogenic oomycetes reveal lifestyle-specific gene expression": Figure S4

a

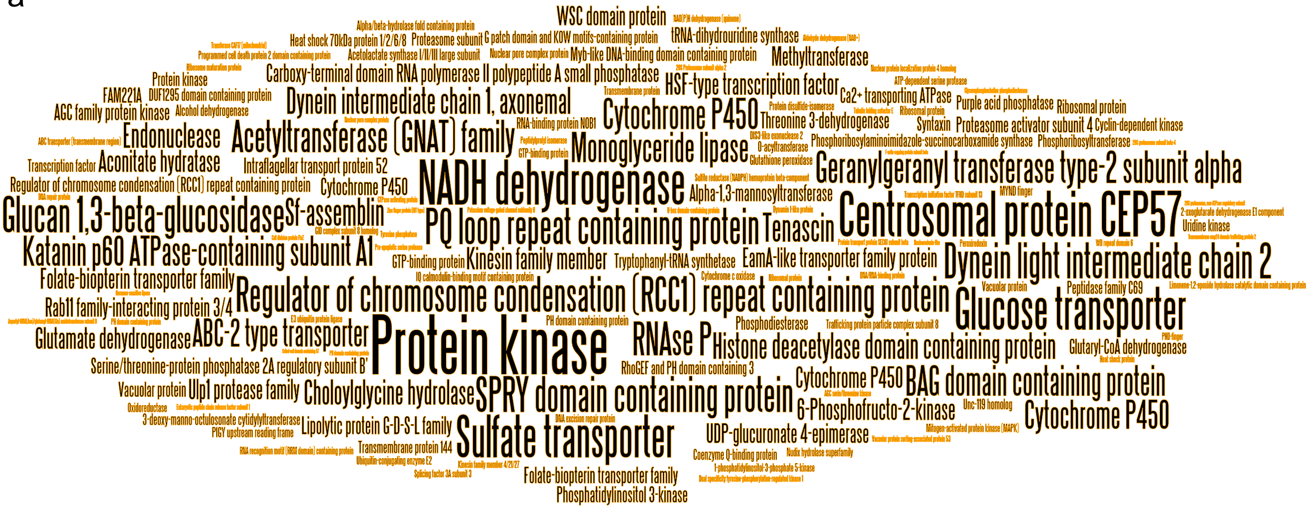


b

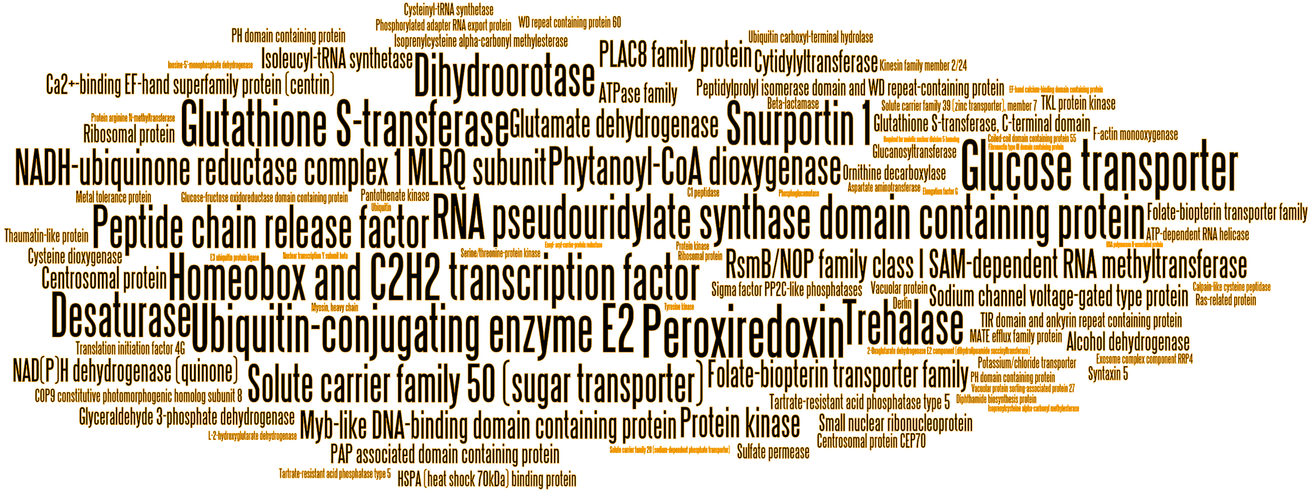


**Figure S4. Annotation of litter-associated genes in clusters C0 and C1.** (a) Wordl of all genes in cluster C0 with an annotation that are significantly up-regulated (FDR ≤ 0.05) in MGLi vs. CMMm. (b) Wordl of all genes that are clustered in cluster C1 and up-regulated (FDR ≤ 0.05) when MGLi is compared to CMMm.
