## Supplementary material for "Comparative analyses of saprotrophy in *Salisapilia sapeloensis* and diverse plant pathogenic oomycetes reveal lifestyle-specific gene expression": Table S1

**Table S1. Transcriptomic and genomic data used in this study for comparative analyses.** Organism indicates which oomycete was used and host denotes which plant was used as host or substrate in the respective studies, tissue indicates the infected or colonized tissue of the plant. Type of data indicates whether we used genomic or transcriptomic data from this organism, reference shows the publication of the transcriptomic or genomic study and from where the data was obtained. Accession and availability give the accession number or location of the data used.

| **Organism** | **Host (if applicable)** | **Tissue/medium** | **Type of data** | **Reference** | **Accession/**  **Availability** |
| --- | --- | --- | --- | --- | --- |
| *Albugo candida* Ac Nc02 | *Arabidopsis thaliana*  MAGIC-107 | leaves | transcriptome | NCBI GenBank; Prince et al. 2017; BMC Biol 15:20 [104] | PRJNA302221 |
| *Albugo laibachii* Al Nc14 | *Arabidopsis thaliana*  MAGIC-107 | leaves | transcriptome | NCBI GenBank; Prince et al. 2017; BMC Biol 15:20 [104] | PRJNA302221 |
| *Hyaloperonspora*  *arabidopsidis* Waco9 | *Arabidopsis thaliana* Col-0 | leaves | transcriptome | NCBI GenBank; Asai et al. 2014; PLoS Pathog 10: e1004443 [21] | PRJNA232536 |
| *Phytophthora*  *infestans* 1306 | *Solanum tuberosum* | tuber, medium (CMA) | transcriptome | Ah-Fong et al. 2017; BMC Genomics 18:764 [23] | Table S2 |
| *Phytophthora parasitica* INRA-310 | *Solanum lycopersicum* | roots | transcriptome | NCBI GenBank | PRJNA168272 |
| *Phytophthora sojae*  pmg(1)-3 | *Glycine max* cv. Williams | hypocotyl | transcriptome | NCBI GenBank; Lin et al. 2014; BMC Genomics 15:18 [103] | PRJNA210431 |
| *Plasmopara halstedii* OS-Ph8-99-BlA4 | *Helianthus annuus* | roots | transcriptome | NCBI GenBank; Sharma et al. 2015; BMC Genomics 16:741 [95] | PRJEB6932 |
| *Pythium ultimum* potato isolate from San Jacinto, CA | *Solanum tuberosum* | tuber, medium (CMA) | transcriptome | Ah-Fong et al. 2017; BMC Genomics 18:764 [23] | Table S2 |
| *S. sapeloensis* | *Spartinia alterniflora* litter | Leaf litter, water associated to leaf litter, medium (CMA) | transcriptome trimmed reads | This study | PRJNA487262 |
| *S. sapeloensis* | *Spartinia alterniflora* litter | Leaf litter, water associated to leaf litter, medium (CMA) | transcriptome *de novo* assembly | This study | PRJNA487262 |
| *Saprolegnia*  *parasitica* CBS 223.65 | Not applicable | Not applicable | genome (protein sequences) | NCBI GenBank;  Jiang et al. 2013; PLoS Genet 9: e1003272 [92] | PRJNA280969PRJNA36583 |
| *Saprolegnia diclina* VS20 | Not applicable | Not applicable | genome (protein sequences) | NCBI GenBank | PRJNA255245PRJNA86859 |
| *Aphanomyces*  *invadans* NJM9701 | Not applicable | Not applicable | genome (protein sequences) | NCBI GenBank | PRJNA258292PRJNA188082 |
| *Aphanomyces astaci* APO3 | Not applicable | Not applicable | genome (protein sequences) | NCBI GenBank | PRJNA264335 PRJNA187372 |
| *Achlya hypogyna* ATTC 48635 | Not applicable | Not applicable | genome (protein sequences) | NCBI GenBank; Misner et al. 2015; Genome Biol Evol 7:120-135 [13] | PRJNA169234 |
| *Thraustotheca*  *clavata* ATCC 34112 | Not applicable | Not applicable | genome (protein sequences) | NCBI GenBank; Misner et al. 2015; Genome Biol Evol 7:120-135 [13] | PRJNA169235 |
| *Pythium irregulare* DAOM BR486 | Not applicable | Not applicable | genome (protein sequences) | Ensembl Protists (Downloads); Adhikari et al. 2013; PLoS One 8: e75072 [91] | pir_scaffolds_v1 |
| *Pythium iwayamai* DAOM BR242034 | Not applicable | Not applicable | genome (protein sequences) | Ensembl Protists (Downloads); Adhikari et al. 2013; PLoS One 8: e75072 [91] | piw_scaffolds_v1 |
| *Pythium ultimum* DAOM BR144 | Not applicable | Not applicable | genome (protein sequences) | Ensembl Protists (Downloads); Lévesque et al. 2010; Genome Biology 11:R73 [89] | pug |
| *Pythium*  *aphanidermatum* DAOM BR444 | Not applicable | Not applicable | genome (protein sequences) | Ensembl Protists (Downloads); Adhikari et al. 2013; PLoS One 8: e75072 [91] | pag1_scaffolds_v1 |
| *Pythium*  *arrhenomanes* ATCC 12531 | Not applicable | Not applicable | genome (protein sequences) | Ensembl Protists (Downloads); Adhikari et al. 2013; PLoS One 8: e75072 [91] | par_scaffolds_v1 |
| *Phytophthora*  *infestans* T30-4 | Not applicable | Not applicable | genome (protein sequences) | NCBI GenBank; Haas et al. 2009; Nature 461: 393–398 [20] | PRJNA49677 PRJNA17665 |
| *Phytophthora*  *parasitica* INRA-310 | Not applicable | Not applicable | genome (protein sequences) | NCBI GenBank | PRJNA259235PRJNA73155 |
| *Phytophthora*  *parasitica* INRA-310 | Not applicable | Not applicable | genome (cDNA sequences) | Ensembl Protists (Downloads) | PP_INRA-310_V2 |
| *Plasmopara halstedii* OS-Ph8-99-BlA4 | Not applicable | Not applicable | genome (protein sequences) | NCBI GenBank; Sharma et al. 2015; BMC Genomics 16:741 [95] | PRJNA314514PRJEB6932 |
| *Plasmopara halstedii* OS-Ph8-99-BlA4 | Not applicable | Not applicable | genome (cDNA sequences) | Ensembl Protists (Downloads); Sharma et al. 2015; BMC Genomics 16:741 [95] | Plasmopara_halstedii_genome |
| *Hylaoperonospora*  *arabidopsidis* Emoy2 | Not applicable | Not applicable | genome (cDNA & protein sequences) | Ensembl Protists (Downloads); Baxter et al. 2010; Science 330:1549-1551 [88] | HyaAraEmoy2_2.0 |
| *Phytophthora*  *lateralis* MPF4 | Not applicable | Not applicable | genome (protein sequences) | Ensembl Protists (Downloads); Quinn et al. 2013; FEMS Microbiol Lett 344:179-185 [93] | MPF4_v1.0 |
| *Phytophthora*  *ramorum* Pr102 | Not applicable | Not applicable | genome (protein sequences) | Ensembl Protists (Downloads); Tyler et al. 2006; Science 313:1261-1266 [87] | ASM14973v1 |
| *Phytophthora sojae* P6497 | Not applicable | Not applicable | genome (protein sequences) | NCBI GenBank; Tyler et al. 2006; Science 313:1261-1266 [87] | PRJNA262907PRJNA17989 |
| *Phytophthora sojae* P6497 | Not applicable | Not applicable | genome (cDNA sequences) | Ensembl Protists (Downloads); Tyler et al. 2006; Science 313:1261-1266 [87] | P_sojae_V3_0 |
| *Phytophthora*  *kernoviae* 00238/432 | Not applicable | Not applicable | genome (protein sequences) | Ensembl Protists (Downloads); Sambles et al. 2015; Genom Data 6:193-194 [94] | PhyKer238_432v1 |
| *Phytopythium vexans* (previously *Pythium vexans*) DAOM BR484 |  |  | genome (protein sequences) | Ensembl Protists (Downloads); Adhikari et al. 2013; PLoS One 8: e75072 [91] | pve_scaffolds_v1 |
| *Albugo laibachiii*  Al Nc14 | Not applicable | Not applicable | genome (cDNA & protein sequences) | Ensembl Protists (Downloads); Kemen et al. 2011; PLoS Biol 9:e1001094 [90] | ENA1 |
| *Albugo candida*  Ac Nc02 | Not applicable | Not applicable | genome (cDNA & protein sequences) | Ensembl Protists (Downloads) | ASM107853v1 |
