## Supplementary material for "Comparative analyses of saprotrophy in *Salisapilia sapeloensis* and diverse plant pathogenic oomycetes reveal lifestyle-specific gene expression": Table S2

| **Gene name** | **Gene ID** | **Primer sequences** | **Annealing temperature [°C]** | **Product length [bp]** |
| --- | --- | --- | --- | --- |
| *CE10* | *Salisap4165_c1_g3* | F: 5’ TGTCTTGGCAGTGTACCGAC 3’  R: 5’ CTCTGCGGAACAAGTGGCTA 3’ | 62 | 178 |
| *GH1* | *Salisap3316_c0_g4* | F: 5’ ACGGGCAAAACCACGAGTAT 3’  R: 5’ CAGCGTCACCTTTTGTTGGG 3’ | 61 | 153 |
| *GH6* | *Salisap3873_c0_g7* | F: 5’ AAGTGCTCTCGGAAGAACCG 3’  R: 5’CGGGTGAAAGTGACGGAGAA 3’ | 61 | 85 |
| *GT57* | *Salisap1930_c0_g1* | F: 5’ TCTCTGCCTTTTGCTGTCGT 3’  R: 5’ ATCTACCGCTTAGCTTCCGC 3’ | 61 | 157 |
| *Myb-related TF* | *Salisap3868_c0_g1* | F: 5’ CGGCACGTCATCCATATCCA 3’  R: 5’ CAAGCTCGTGGACAAAACGG 3’ | 62 | 193 |
| *NF-YB TF* | *Salisap4629_c0_g1* | F: 5’ CGAAATGAACTCGGACACGC 3’  R: 5’ GCAGATGAGATTCGGGAGCA 3’ | 62 | 141 |
| *H2A* | *Salisap2927_c0_g1* | F: 5’ GGAGGCATGGAGAAACGTCA 3’  R: 5’ ATCATCCCGCGTCACATTCA 3’ | 61 | 226 |

**Table S2. Primers used in the qRT-PCRs.** Gene name and gene ID are given in the correspondingly named column. Primer sequences for forward and reverse primer are given for each tested gene. F indicates the forward primer and R indicates the reverse primer. The annealing temperature used for the qRT-PCR reaction is given in the column ‘Annealing temperature [°C] and the base pair [bp] length of the PCR product is given in the respective column.
